# Promiscuous Metal Site in Hepatitis B Virus X Protein Binds an Fe-S Cluster

**DOI:** 10.64898/2026.01.20.700672

**Authors:** Jiahua Chen, Michelle Langton, Patrick Cao, Avital Aaron, Jackson Ho, Eranthie Weerapana, Deborah L. Perlstein, Alexey Silakov, Daniel W. Bak, Maria-Eirini Pandelia

## Abstract

The Hepatitis B virus (HBV) regulatory protein HBx is essential for viral replication and pathogenesis, yet its cofactor specificity and ligand environment remain poorly defined. Although HBx binds either an Fe-S cluster or Zn, its intrinsic disorder and mutational tolerance have hindered its precise characterization. Here, we integrate chemoproteomics with HYSCORE spectroscopy to identify the metal-coordinating ligands in HBx. Histidine coordination is excluded, while C61, C69, C143, and C148 emerge as primary cysteine ligands for the Fe-S cluster, with C137 acting as a conditional ligand. These residues also bind Zn and are associated with HBx transactivation and clinically relevant variants. HBx engages the host cytosolic Fe-S machinery and displays sensitivity to Fe-S-targeting reagents, behavior consistent with Fe-S cluster acquisition and lability. Together, these findings suggest that HBx functionally behaves as an Fe-S cluster-associated protein, highlighting a potentially druggable vulnerability in HBV replication.

**Significance Statement:** Fe-S clusters are emerging as key cofactors in viral replication but are often mistaken for Zn due to O_2_-sensitivity and shared cysteine coordination. The Hepatitis B virus HBx protein, essential for viral replication and hepatocarcinogenesis, has long been mechanistically intractable, with debate over its metallocofactor. Here, we provide evidence that HBx coordinates an Fe-S cluster, placing it within the growing family of viral Fe-S-cluster-containing proteins. Using chemoproteomics, we identify its cysteine ligands, overcoming limitations of mutational analysis in disordered proteins. Although HBx binds both an Fe-S cluster and Zn, its interaction with human Fe-S assembly factors suggests a functional link to Fe-S cluster biology, while its sensitivity to TEMPOL and NO reveals a potentially druggable vulnerabilityin HBx.

## Introduction

Chronic Hepatitis B virus (HBV) infection affects approximately 300 million people worldwide and can lead to life-threatening liver diseases, including cirrhosis and hepatocellular carcinoma (1, 2). At the core of HBV infectivity lies the virus-encoded X protein (HBx). This 17 kDa multifunctional oncoprotein is required for viral replication and interferes with numerous cellular signaling pathways, contributing to the development of HBV-associated liver diseases (3–5). Recent studies have shown that HBx is a *bona fide* metalloprotein capable of coordinating an iron-sulfur (Fe-S) cluster or a Zn ion (6–10). These metallocofactors are essential for many proteins across all domains of life, including viruses, where they fulfill diverse structural and functional roles (7, 11–14). However, for HBx, the precise *modus operandum* of the Fe-S cluster or Zn has been little explored, and this cofactor plurality is of unknown functional significance.

Both Fe-S clusters and Zn can be tetrahedrally coordinated by cysteine and histidine residues arranged in CCCC, CCCH, and CCHH motifs (7, 15, 16). This shared coordination geometry allows the two metallocofactors to replace each other at the same binding site, either in a functionally interchangeable or exclusive manner. However, due to the inherent O_2_-sensitivity of Fe-S clusters and the use of nonspecific metal-containing media during protein expression, Zn incorporation into these motifs is strongly biased (7, 15). As a result, these motifs are frequently referred to as zinc-finger (ZF) motifs, a term that can be misleading when based solely on the primary sequence. This misnomer, along with the binding promiscuity and instability of Fe-S cofactors, has often led to Fe-S cluster proteins being overlooked or misannotated as Zn-binding proteins (7, 14, 15). Some well-studied cases include ferredoxin Fdx and the Fe-S cluster donor proteins IscA and IscU, which contain ZF motifs and bind Zn with high affinity even when isolated under O_2_-free conditions (17–19). To enable Fe-S cluster incorporation in these proteins, Zn needs to be removed. In some proteins that contain multiple ZF domains, Zn and Fe-S cluster binding can be functionally codependent. The cleavage and polyadenylation specificity factor 30 (CPSF30) possesses five non-metal specific CCCH motifs, with high-affinity RNA binding requiring the simultaneous coordination of at least one [2Fe-2S] cluster and two Zn ions (15, 20). A similar synergistic effect has been observed in the SARS-CoV-2 Nsp13 helicase, which harbors three ZF motifs (21). These examples highlight the need to define the cofactor specificity and metal-binding ligands in HBx, whose ability to bind both Fe-S cluster and Zn poses similar challenges to both its annotation and functional understanding.

Fe-S clusters are increasingly recognized as important functional elements in viral proteins that govern viral replication, making them attractive drug targets for suppressing infections (7, 14). In Nsp12, the catalytic subunit of the SARS-CoV-2 RNA-dependent RNA polymerase, the coordination of a [4Fe-4S] cluster is essential for polymerase activity and can be disrupted by nitroxides, offering a valuable alternative antiviral avenue (14, 22–24). More recently, [4Fe-4S] clusters were identified in both Nsp10 and Nsp14, which enhance the methyltransferase activities (24). The Merkel cell polyomavirus small T antigen was initially thought to bind two Zn ions but was later found to coordinate a [2Fe-2S] and a [4Fe-4S] cluster, which are crucial for viral replication (25). Furthermore, a [2Fe-2S] cluster at the interface of the rotavirus NSP5 dimer has been shown to be essential for single-stranded RNA binding (26). Fe-S clusters have also been discovered in the small protein R633b from the Megavirinae giant viruses, albeit their functional roles are yet to be established (27).

HBx, like several other nonstructural viral proteins, displays a dual metal-binding behavior, associating either with a redox-active Fe-S cluster, an O_2_-sensitive [4Fe-4S] species that converts to a stable [2Fe-2S] form under ambient conditions, or a redox-inert Zn (6, 8, 10). Although both cofactors can serve structural roles, their distinct chemical properties, particularly the redox sensitivity of Fe, confer Fe-S clusters additional roles in oxygen-sensing, degradation signaling, or catalysis (11, 28). Whether Fe-S clusters and Zn function interchangeably at the same site in HBx remains unclear. However, growing evidence supports Fe-S clusters as genuine viral cofactors, suggesting that at least one is functionally relevant in HBx (14, 22). A common strategy to resolve such ambiguity is to determine whether the viral protein engages with the host Fe-S cluster assembly pathways, such as the Iron-Sulfur cluster (ISC) or cytosolic iron sulfur cluster assembly (CIA) machinery. Because viruses lack the ability to synthesize Fe-S clusters *de novo* any Fe-S cluster-dependent viral protein must rely on host pathways for cofactor acquisition (14, 29). Establishing whether HBx interacts with these host assembly pathways, therefore, represents a critical step in defining the chemical identity of its metallocofactor. Beyond clarifying the HBx metal-dependence, this information is important for advancing our understanding of virus-host interactions and for opening new therapeutic avenues to target HBx in HBV infection.

HBx from different HBV genotypes contains multiple conserved cysteines and a single conserved histidine, which are potential ligands for either an Fe-S cluster or Zn (6, 8, 10, 15). The conserved cysteines have been frequently identified as functional hotspots in HBx, with numerous studies showing that their substitution disrupts key HBx-mediated processes, including transcriptional activation, subcellular localization, and protein-protein interactions (4, 30–32). Although the molecular basis for these effects is poorly understood, the involvement of these cysteines in metallocofactor coordination suggests that Fe-S cluster or Zn may play important regulatory roles in HBx function. A crucial step toward elucidating HBx-mediated mechanisms is therefore to define the relationship between metallocofactor identity and the metal-binding ligands with conserved residues, whose perturbation impairs HBx activity. Zn has been previously proposed to be coordinated via a CCCH motif comprising C61, C69, C137, and H139 (10). While these residues are individually important for the targeted degradation of the host restriction factor chromosome 5/6 (Smc5/6), a key antiviral defense against HBV, direct evidence linking Zn coordination to HBx function has not been made (10). Furthermore, there is no definitive support for H139 as a Zn ligand, and substitution of the entire CCCH motif does not abolish Zn binding in HBx (10).

In this study, we employed the well-characterized recombinant HBx to systematically investigate the metal-binding properties of HBx and its functional significance. Specifically, we (i) identified the candidate ligands to the Fe-S cluster, (ii) evaluated the specificity and overlap in ligation patterns between the Fe-S cluster and Zn, (iii) demonstrated interaction of HBx with the host Fe-S biosynthesis factors CIAO1 and CIAO2B, and (iv) screened cell-permeable small molecules for their ability to disrupt the HBx Fe-S cluster. Collectively, these findings are consistent with the identification of HBx as an Fe-S cluster-dependent protein and provide a mechanistic framework linking metal coordination in HBx to HBV pathogenesis.

## Results

### Computational prediction of metal-binding ligands

HBx proteins from all ten HBV genotypes (A-J) share more than 80% sequence identity and contain eight strictly conserved cysteine residues (C7, C17, C61, C69, C115, C137, C143, and C148) (**Figs. 1A and S1**), some of which are proposed to coordinate either an Fe-S cluster or Zn (6, 10). Additionally, some genotypes contain extra cysteines at positions 6 (B-J), 26 (E and H), and 78 (A and J). A strictly conserved histidine at position 139 is also thought to act as a metallocofactor ligand (**Figs. 1A and S1**) (10). In a first step, we employed AlphaFold to gain structural insights into the spatial arrangement of these conserved residues (33). H139 and the majority of the cysteines (C61, C69, C78, C137, C143, and C148) are predicted to localize in an unstructured region, while C7 and C17 are situated within an antiparallel β-sheet (residues 5-10; 15-20), and C115 is positioned in the middle of an α-helix (residues 105-126) (**Fig. 1A**) (9). The observed clustering of C61, C69, C137, H139, C143, and C148 suggests a possible metal-binding pocket. However, the high degree of apparent disorder, combined with the low-confidence AlphaFold predictions, makes it particularly difficult to reliably pinpoint the metal-binding site(s) in HBx. Most residues have pLDDT scores below 50, regardless of whether Zn is included in the modeling (Fe-S cluster modeling remains unfeasible), making it difficult to confidently identify the metal ligands using this approach (33).

**Figure 1.**
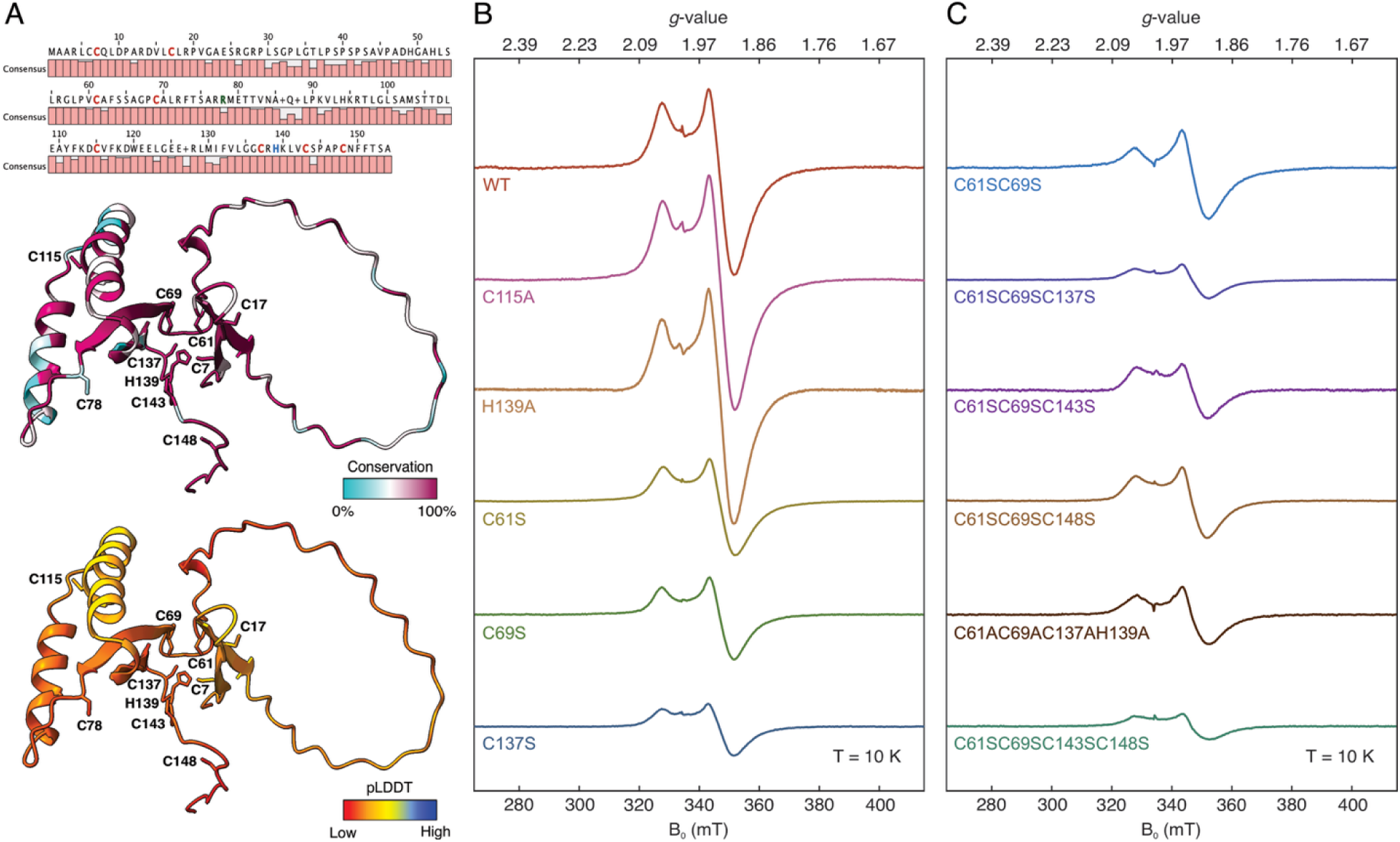
Consensus sequence, structural models, and X-band CW EPR spectra of WT and variant HBx. **A**. (top) Consensus sequence plot of HBx from the ten HBV genotypes, A to J. The eight cysteines and one histidine that are strictly conserved across the 10 genotypes are shown in red and blue, respectively. R78, which aligns with a conserved cysteine residue in HBV genotypes A and J, is shown in green. (middle) HBx (genotype A2) AlphaFold model colored according to sequence conservation of the 10 representative sequences from genotypes A to J. (bottom) HBx AlphaFold model colored according to the predicted local-distance difference test (pLDDT) score. pLDDT is below 60 for all residues. **B**. and **C**. CW EPR spectra of the sodium dithionite-reduced WT and variant DsbC-HBx monitoring assembly of [4Fe-4S]^1+^ clusters. WT and variant DsbC-HBx were co-expressed with the pDB1282 plasmid in M9 media supplemented with Fe(NH_4_)_2_(SO_4_)_2_, then isolated under O_2_-free conditions and chemically reconstituted during lysis. The intensities have been normalized to account for the same protein concentration. Experimental conditions: microwave frequency = 9.38 GHz, temperature = 10 K, microwave power = 0.64 mW, modulation amplitude = 1mT.

### Cysteine substitutions do not abolish Fe-S cluster coordination

Given the limited reliability of computational predictions and the intrinsic disorder nature of HBx, we employed a systematic mutational approach to map residues involved in Fe-S cluster coordination by generating a series of single- and multiple-point substitutions targeting residues that are either functionally important or previously reported to be involved in metal binding (9, 10, 30–32, 34–36). Alanine and serine substitutions produced indistinguishable EPR spectra in terms of both lineshape and g-values, indicating they can be used interchangeably and that serine cannot serve as a cluster ligand (37). For these experiments, we used our previously well-established DsbC-HBx construct that was anaerobically isolated and reconstituted during lysis to afford homogeneous enrichment of a [4Fe-4S]^2+^ cofactor (6). To generate the paramagnetic [4Fe-4S]^1+^ form (indicative of Fe-S cluster incorporation), the WT and variant DsbC-HBx were reduced with sodium dithionite, and the resulting CW EPR spectra are shown in **Figure 1**.

WT DsbC-HBx exhibits an EPR signal of axial symmetry and principal *g*-values of 2.04 and 1.94, reminiscent of low-spin (*S* = 1/2) [4Fe-4S]^1+^ clusters (**Fig. 1B**). Single-point substitutions of the mitochondrial targeting residue C115 (C115A) and the proposed metal binding residue H139 (H139A) exhibited WT-like spectra in terms of lineshape, *g*-values, and intensity. In contrast, single-point substitutions of the cysteines C61S, C69S, and C137S that are within the CCCH motif previously proposed to coordinate Zn, resulted in a significant reduction in [4Fe-4S]^1+^ cluster intensity to 56%, 39%, and 26%, respectively, relative to the WT protein (**Figs. 1B and S2**). The pronounced loss of cluster signal in the C137S variant may be partially attributed to reduced overall protein stability compared with C61S and C69S variants, potentially further compromising proper cluster incorporation. More extensive substitutions within the CCCH motif, such as C61SC69S, C61SC69SC137S, and C61AC69AC137AH139A, appreciably reduced Fe-S cluster incorporation to varying extents; however, none of those CCCH motif variants completely abolished Fe-S cluster formation (**Fig. 1C**).

We also introduced substitutions at the HBx C-terminus, a region that is essential for protein stability, transcriptional activation, and HBV replication (35, 38, 39). Similar to substitutions at the proposed CCCH motif, serine or alanine replacements of C-terminal cysteines, regardless of their extent and number, did not eliminate Fe-S cluster incorporation. The triple variants, C61SC69SC143S and C61SC69SC148S, retained approximately 40% of [4Fe-4S]^1+^ cluster intensity, while the quadruple variant C61SC69SC143SC148S resulted in the lowest Fe-S cluster incorporation amounting to 16% of that of the WT protein (**Figs. 1C and S2)**. Other C-terminal variants, such as C61AC69AC115AC137A and C137AC143AC148A, displayed 30% and 37% yield in a [4Fe-4S]^1+^ cluster signal, respectively (**Fig. S3**). More extensive substitutions of cysteines at the negative transcriptional regulatory N-terminus (i.e., C7AC17AC61AC69AC78AC115A) still showed an appreciable [4Fe-4S] cluster signal (**Fig. S3**). Complete abolishment of the Fe-S cluster signal was only achieved in a variant in which all cysteines were replaced by alanines (**Fig. S3**). These findings demonstrate a certain plasticity in cofactor coordination, most likely facilitated by the apparent conformational disorder of HBx, as predicted by AlphaFold and supported by CD and NMR studies (33, 40, 41). The results of mutational scanning were comparable whether monitoring the incorporation of a [4Fe-4S] cluster under anaerobic conditions or a [2Fe-2S] cluster under ambient conditions, demonstrating that both cluster forms behave similarly in solution (**Fig. S4**).

### H139 is not an Fe-S cluster ligand

To unambiguously determine whether H139 is involved in Fe-S cluster ligation, HYSCORE experiments were carried out on the WT and H139A DsbC-HBx (**Figs. 2 and S5**). The HYSCORE spectra of the anaerobically isolated and dithionite-reduced proteins were collected at a magnetic field of 345 mT, corresponding to the principal value *g* of 1.94 of the EPR signal of the paramagnetic [4Fe-4S]^1+^ cluster. The spectrum of the WT protein contains peaks along the diagonal that are representative of matrix ^14^N nitrogens (4 MHz) and ^1^H protons (15 MHz). No off-diagonal signals due to strongly coupled ^14^N nuclei were observed, indicating the absence of an imidazole nitrogen as a ligand to the [4Fe-4S]^1+^ center (42). The HYSCORE spectrum of the H139A variant was identical to that of the WT, further confirming that the conserved H139 does not serve as a ligand to the Fe-S cluster (**Fig. 2**). These experiments were also performed with samples that were cryo-reduced, a method that would minimize any disorder introduced by chemical reduction of the Fe-S cluster, with identical results. (**Fig. S6**) These findings align well with the results from the CW EPR experiments, in which the WT and H139A variant exhibited identical lineshapes and *g*-values, as well as a similar extent of Fe-S cluster incorporation (**Figs. 1 and S5**).

**Figure 2.**
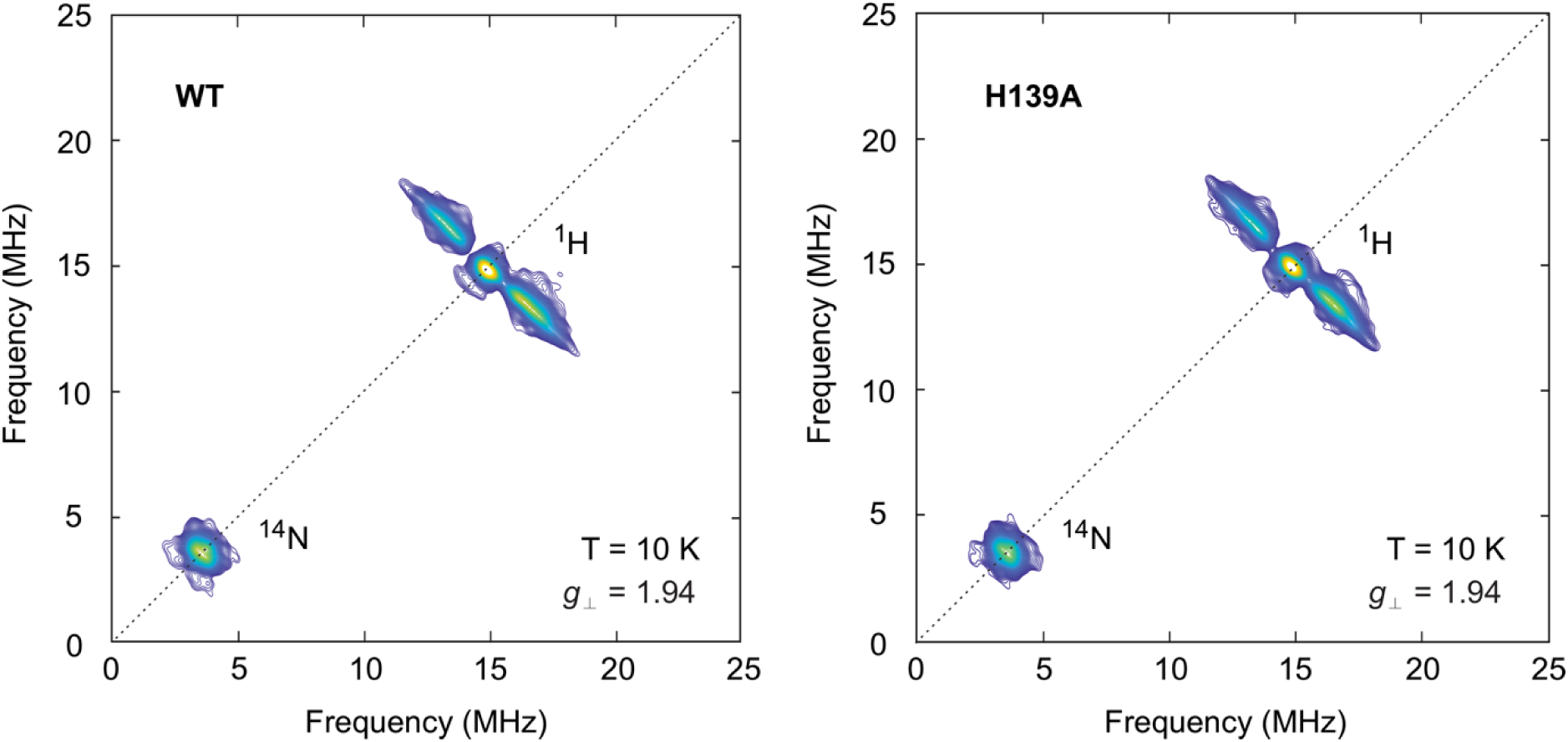
X-band HYSCORE spectra of the WT and H139A DsbC-HBx. The anaerobically isolated WT (left) and H139A (right) DsbC-HBx were the pDB1282 plasmid in M9 media supplementated with Fe(NH_4_)_2_(SO_4_)_2_, then chemically reconstituted during lysis, and then the purified proteins with [4Fe-4S]^2+^ form were reduced with an excess of sodium dithionite (6 mM) for 20 min to obtain the Fe-S cluster in the [4Fe-4S]^1+^ form. The HYCORE spectra have been recorded at a magnetic field of 345 mT, corresponding to the principal value *g*_⊥_ of 1.94 (**Figs. 1 and S5**). Experimental conditions: microwave frequency = 9.44 GHz, temperature 10 K, *τ* = 132 ns, *t*(π/2) = 8 ns.

### Competitive Binding of an Fe-S cluster and Zn in HBx

Because EPR mutational scanning revealed similar solution behavior and cysteine substitution effects for [4Fe-4S]- and [2Fe-2S]-containing HBx, we proceeded to quantify the extent of cofactor incorporation by ICP-AES using aerobically isolated protein. The WT DsbC-HBx protein was expressed in LB media supplemented with Fe and/or Zn to examine the impact of metal availability on the extent of Fe-S cluster or Zn incorporation (**Fig. 3**). The Fe content directly reports on [2Fe-2S] incorporation, as confirmed by optical and Mössbauer studies, and can be converted to cluster equivalents if divided by 2 (6). For the WT protein, under all conditions examined, the molar amounts of the [2Fe-2S] cluster and Zn per protein are consistently less than one. In the absence of exogenous metals, the extent of [2Fe-2S] and Zn incorporation is roughly equivalent to approximately 0.4 mol cofactor per mol protein. Considering that the reported concentrations of Fe and Zn in *E. coli* cells grown in LB are comparable and in the low millimolar range, HBx appears to exhibit similar affinities for both a [2Fe-2S] cluster and Zn (43). Addition of Fe to the LB media results in a 1.6-fold increase in Fe-S cluster binding and a 1.4-fold decrease in Zn incorporation (**Fig. 3**), demonstrating a competition between these two cofactors and alluding to the presence of common ligand binding site(s). Addition of Zn to the LB media has an analogous effect, with a 2.3-fold decrease in Fe-S cluster binding and a 2.2-fold increase in Zn incorporation. When Fe and Zn are simultaneously added in equimolar amounts, incorporation of the [2Fe-2S] cluster and Zn exhibits a 1:2 ratio. The apparent compromise in Fe-S cluster content when both metals are added may stem from competitive Zn binding in HBx and/or Zn inhibition of the Fe-S cluster biogenesis machinery, as these experiments involve co-expression with the pDB1282 plasmid that encodes the isc operon, the core Fe-S cluster assembly pathway (6, 17, 18, 44, 45). Notably, Fe-S cluster incorporation was never completely abolished under any Zn enrichment condition, a property shared with other Fe-S cluster binding proteins, such as Fdx and IscA (18).

**Figure 3.**
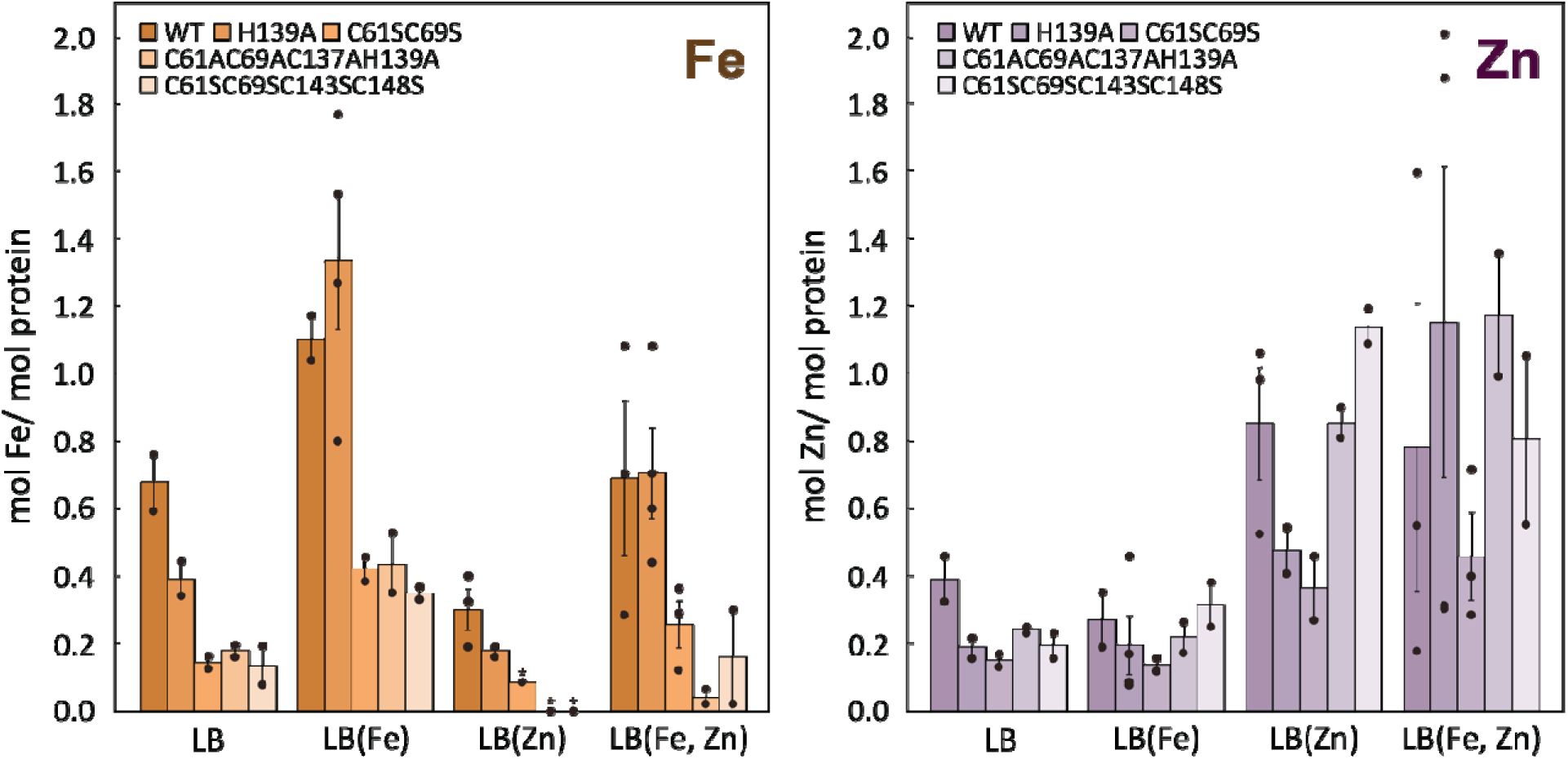
Metal binding of WT and variant DsbC-HBx. Elemental analysis of Fe and Zn content for aerobically isolated WT and variant DsbC-HBx. Proteins were expressed with the pDB1282 plasmid under various metal supplementation conditions: LB only, LB with 250 μM Fe(NH_4_)_2_(SO_4_)_2_, LB with 250 μM ZnSO_4_, and LB with 250 μM each Fe(NH_4_)_2_(SO_4_)_2_ and ZnSO_4_. Individual data points are shown as circles. Circles marked with an asterisk represent data from two independent expressions and purifications that yielded identical results. Error bars indicate the standard error of the mean.

### C61 and C69 are common ligands to the Fe-S cluster and Zn

To further define the potential cofactor ligands and identify any shared binding sites, we systematically examined Fe-S cluster and Zn incorporation across DsbC-HBx variants under varying supplementation conditions (**Fig. 3**). The H139A variant exhibited comparable Fe-S cluster incorporation to the WT protein under all tested conditions, consistent with our EPR and HYSCORE data (**Figs. 1B and 2**), and confirming that H139 does not coordinate the Fe-S cluster. In contrast, Zn incorporation in the H139 variant was highly variable, remaining unchanged in some conditions (e.g., LB(Fe, Zn)) but reduced in others (e.g., LB(Zn)) relative to the WT protein, rendering its role in Zn coordination inconclusive. In comparison, the C61SC69S double variant, which exhibits an appreciable reduction in the EPR Fe-S cluster signal intensity (**Figs. 1C and S4**), displayed substantially diminished binding of both cofactors. Specifically, substitution of C61 and C69 with a serine led to at least a 2.6-fold and 1.7-fold decrease in Fe-S cluster and Zn binding, respectively, when compared to the WT protein, suggesting that these two cysteines serve as common ligands for both metallocofactors.

We next assessed Fe-S cluster and Zn incorporation in the more extensively substituted quadruple variants that include C61 and C69. The C61AC69AC137AH139A variant, which eliminates the proposed CCCH metal-binding motif, showed reduced Fe-S cluster incorporation under all tested conditions compared to the WT protein. Similarly, the C61SC69SC143SC148S variant, which showed the lowest Fe-S cluster incorporation by EPR, exhibited a substantial loss in Fe-S cluster binding compared to the WT protein (**Figs. 1C and S2**). These findings suggest that increasing the number of cysteine substitutions progressively impairs Fe-S cluster incorporation. In contrast, the effects of these mutations on Zn binding were less straightforward. Despite the extensive substitutions, both quadruple variants retained Zn levels comparable to or even exceeding those of WT and notably higher than the C61SC69S double variant. This unexpected result may reflect altered protein conformations that permit nonspecific or adventitious Zn binding. Together, elemental analysis and mutagenesis studies support the involvement of specific residues in Fe-S cluster and Zn binding. However, pinpointing definitive ligands remains challenging, likely due to the structural flexibility and disorder of the HBx polypeptide.

### C61, C69, C143, and C148 are key Fe-S cluster ligands

To more precisely identify the cysteine residues involved in the Fe-S cluster coordination, we employed a chemoproteomic strategy that quantifies the differential reactivity of cysteines in both the metal-free and metal-bound states (46, 47). Apo forms of WT and variant DsbC-HBx were expressed in M9 media devoid of any metal ions and isolated under anaerobic conditions. These apo forms were then subjected to an *in vitro* reconstitution using the cysteine desulfurase IscS, either in the presence of iron and L-cysteine to generate holo-HBx enriched in [4Fe-4S]^2+^ clusters or in its absence to produce apo-HBx (**Fig. 4A**). Apo- and holo-HBx were then differentially labeled with isotopically distinct forms of the thiol alkylating agent NEM, using NEM-d_0_ (light, L) for apo-HBx and NEM-d_5_ (heavy, H) for holo-HBx. Following proteolytic digestion, the NEM-labeled peptides were analyzed and quantified by LC-MS/MS. Fe-S cluster coordination reduces the accessibility and reactivity of cysteines, resulting in decreased NEM labeling in the holoprotein relative to the apoprotein. Thus, the cysteines involved in cluster binding are expected to show L/H ratios greater than 1, with higher L/H ratios indicating greater protection from alkylation due to Fe-S cluster coordination (**Fig. 4 and Table S1-S3**).

**Figure 4.**
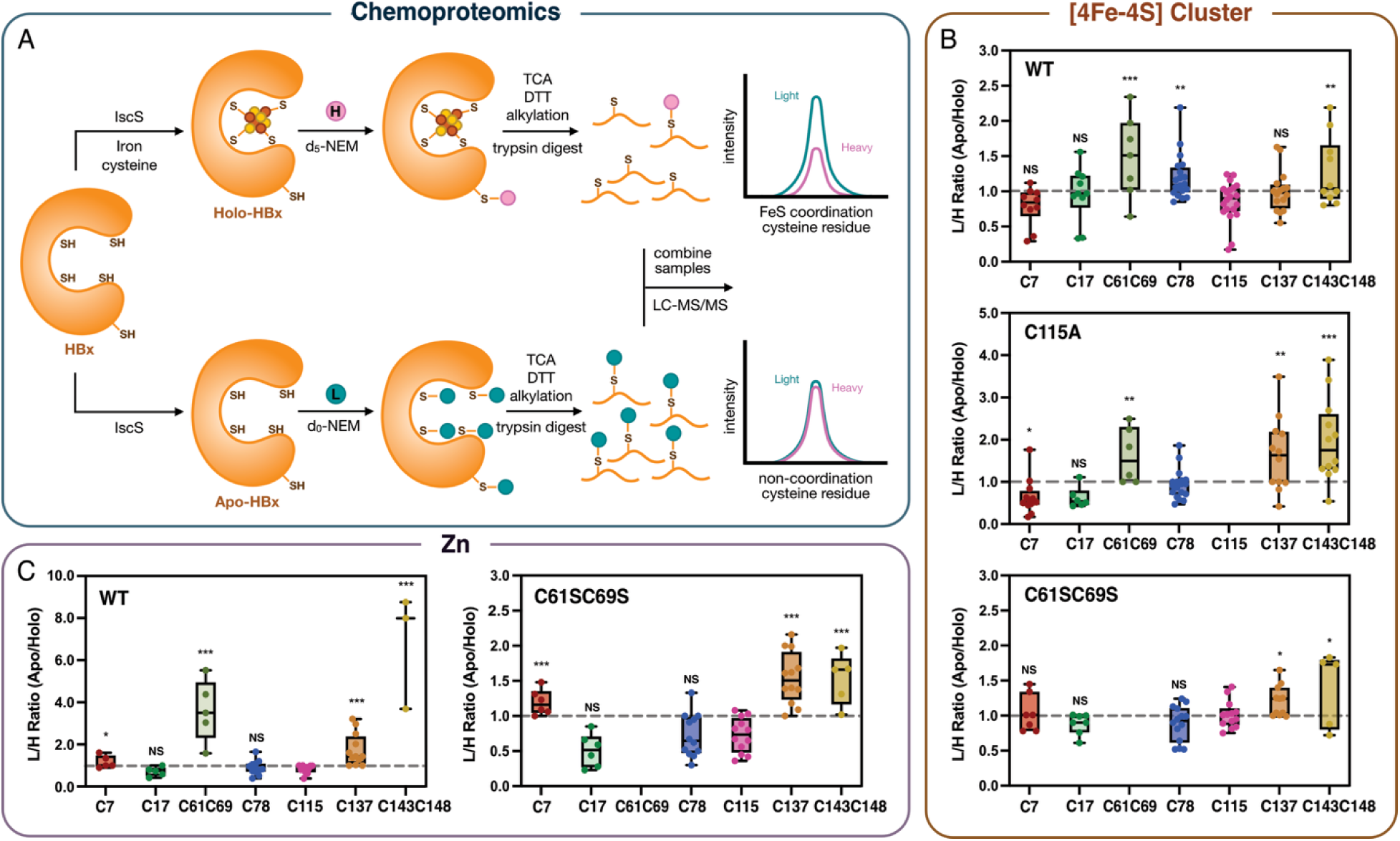
Chemoproteomics pinpoints cysteine ligands involved in Fe-S cluster and Zn coordination in HBx. **A.** Schematic representation of the workflow for isotopic cysteine labeling and LC-MS/MS analysis. The DsbC-HBx was expressed in the M9 media without metal supplemetattion and the anaerobically isolated protein was (**B**) IscS-mediated reconstituted with a [4Fe-4S] cluster and (**C**) chemically reconstituted with Zn. The cysteine reactivity (light:heavy, L/H) ratios are displayed as box plots. For each cysteine residue, each dot represents an independent L/H ratio measurement. The median L/H ratio value across all cysteine-containing peptides (C7, C17, C61C69, C78, C115, C137, and C143C148) is indicated by a dashed line. Statistical significance is determined by the independent *t*-test analysis and is shown as NS (not significant), *\*p* < 0.05, *\*\*p* < 0.01, and *\*\*\*p* < 0.001.

Analysis of the NEM reactivity profiles showed that in the WT protein, the C61, C69-containing peptide exhibited a significantly elevated average L/H ratio of 1.49 (*p*□<□0.001, compared to the C115-containing peptide), consistent with cluster coordination by one or both cysteines (**Fig. 4B and Table S1**). Although individual contributions of C61 and C69 could not be resolved due to their presence on the same tryptic peptide, these findings align with EPR and elemental analyses, supporting the involvement of both residues in cluster coordination (**Figs. 1 and 3**). Similarly, peptides containing C78, and C143, C148 also displayed average L/H ratios greater than 1, suggesting their involvement in cluster coordination. In contrast, C7, C17, C115, and C137 exhibited L/H ratios of approximately 1, indicating minimal or no involvement in Fe-S cluster binding. In the C115A variant, which retains a comparable Fe-S content to that of the WT, C17 and C7 retained L/H ratios below 1, confirming that these residues do not contribute to cluster coordination. The C61, C69 peptide again exhibited elevated L/H ratios, while the C143, C148 peptide displayed an even higher average L/H ratio of 1.97 (*p*□<□0.001). Although the general trend is conserved, the C115A substitution alters the L/H ratios of C78 and C137 compared to the WT protein. C78 displayed an average L/H ratio of 0.94, indicating a lower likelihood of being a cluster ligand, whereas C137 showed an average L/H ratio of 1.64 (*p*□<□0.01), marking it as a possible cluster ligand. Despite some variability in the L/H ratios corresponding to C137, there is a good overall agreement with the EPR and elemental analysis, supporting the involvement of C61, C69, C137, C143, and C148 in Fe-S cluster coordination (**Figs. 1B, 1C, and 3**).

Given the high structural plasticity of HBx, which has deterred any solid attempts to uniquely identify the Fe-S cluster ligands through single or multiple cysteine substitutions, we extended our chemoproteomic analysis to the C61SC69S variant to probe which cysteines could act as ligands in the absence of these residues (**Fig. 4B and Table S1**). In this context, C7, C17, C78, and C115 again showed L/H ratios near 1, whereas peptides containing C143, C148 and C137 exhibited elevated L/H ratios of 1.39 and 1.21, respectively (*p*□<□0.05). These data suggest that in the absence of C61 and C69, C137 can act as an auxiliary ligand, with C143 and C148 consistently contributing to cluster coordination.

To confirm that these observations were not an artifact resulting from the sample preparation method, we conducted parallel experiments using independently expressed and purified apo- and holo-HBx (**Fig. S7**). This approach also enabled us to additionally investigate whether the [2Fe-2S] and [4Fe-4S] clusters share a common ligation site, as proposed in our previous spectroscopic studies (6). The C61, C69 and C143, C148 peptides consistently exhibited the highest L/H ratios across both cluster types, indicating that the same ligand set coordinates both Fe-S cluster forms (**Fig. S7 and Table S2**). Collectively, these results support a model in which Fe-S cluster binding is primarily mediated by C61, C69, C143, and C148, with C137 serving as a potential auxiliary ligand. While the exact coordination environment could not be unambiguously assigned, our findings identify the most probable and biologically relevant residues involved in Fe-S cluster assembly, providing a foundation for further functional investigations into the metal-dependent regulation of HBx.

### Fe-S Cluster and Zn share a common binding site

We further extended the chemoproteomics approach to map how Zn is coordinated in HBx (48, 49). The holoprotein was generated by chemically reconstituting anaerobically isolated apo DsbC-HBx with a stoichiometric amount of Zn. In the Zn-enriched WT protein, cysteines C17, C78, and C115 displayed average L/H ratios near 1, indicating that these residues are unlikely to participate in Zn binding (**Fig. 4C and Table S3**). In contrast, the C61, C69 and C143, C148 peptides showed significantly elevated L/H ratios (3.61, *p*□<□0.001) and (6.81, *p*□<□0.001), respectively, consistent with their roles in Fe-S cluster coordination and suggesting they also contribute to Zn binding. C7 and C137 displayed modest by statistically significant increases in L/H ratios of 1.17 (p□<□0.05) and 1.74 (*p*□<□0.001), respectively, indicating possible auxiliary involvement in Zn coordination. This overall pattern of L/H ratios was preserved in the C61SC69S variant. In the double variant, the C143, C148 peptide retained elevated L/H ratios, albeit lower than in the WT, while C7 and C137 maintained L/H ratios similar to those observed in WT HBx. These data suggest that Zn and Fe-S clusters are coordinated by a shared set of cysteine residues, primarily C61, C69, C143, and C148. C137 may serve as an auxiliary ligand for both metallocofactors, while C7 appears to contribute more selectively to Zn binding. Together, these findings underscore the flexible and overlapping nature of metal coordination in HBx, likely reflecting its intrinsically disordered nature.

### HBx interacts with the cytosolic Fe-S cluster assembly proteins CIAO1 and CIAO2B

The ability of HBx to bind both an Fe-S cluster and Zn does not reveal which of the two metallocofactors is biologically relevant. To determine whether HBx is a *bona fide* Fe-S cluster binding protein, we examined whether HBx engages with components of the host Fe-S cluster assembly machinery, as viruses are unable to synthesize these cofactors *de novo* (14). Given the nucleocytoplasmic localization of HBx, we focused on the cytosolic Fe-S cluster targeting complex (CTC), specifically investigating potential interactions with the human CTC components, CIAO1 and CIAO2B (50).

We first evaluated whether HBx interacts with CIAO1 by co-expressing His-tagged HBx and His-Strep-tagged CIAO1 *in E. coli,* followed by strep-tactin affinity purification. This approach alleviates possible interference of the larger DsbC fusion tag with protein-protein interactions and enables evaluation of whether CIAO1 affects HBx solubility. When expressed alone, HBx was insoluble under the tested conditions (**Fig. S8**); however, co-expression with CIAO1 improved its solubility and resulted in its copurification in the elution fraction, consistent with a direct interaction between the two proteins (**Figs. 5A and S9A**). Because *E. coli* lacks the CIA system, these results cannot be attributed to bridging interactions mediated by other CIA factors, as can occur in pull-down assays with eukaryotic hosts. We next examined whether HBx could also associate with CIAO1 in the presence of CIAO2B, another core CTC component. His- and GST-tagged CIAO2B was co-expressed with CIAO1 alone or with both CIAO1 and HBx, followed by GSTrap purification using CIAO2B as bait. As expected, CIAO1 was detected in the elution irrespective of HBx, consistent with the established CIAO1:CIAO2B interaction (**Figs. 5B and S9B**). Importantly, HBx was detected in the elution when both CIAO1 and CIAO2B were co-expressed, indicating the formation of a ternary CIAO1:CIAO2B:HBx complex. Protein identities were confirmed by LC-MS/MS (**Table S4**). The weaker HBx band intensity relative to CIAO1 and CIAO2B may reflect its limited solubility, lower binding affinity, or the absence of additional stabilizing CTC components (e.g., MMS19) under our experimental conditions. Nevertheless, these data suggest that HBx engages the complete CTC, analogous to other client proteins that acquire Fe-S clusters through this pathway.

**Figure 5.**
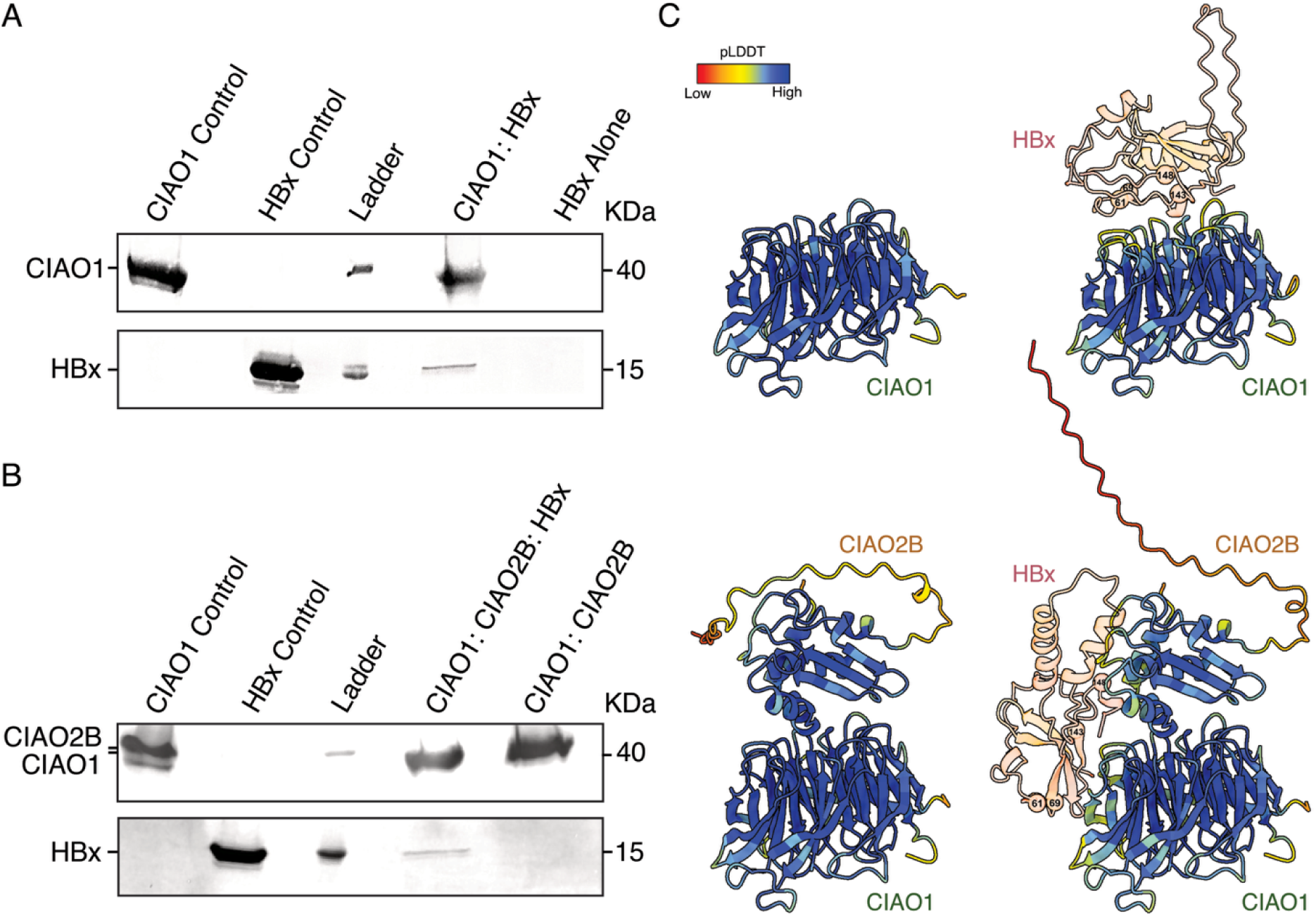
HBx interacts with the human cytosolic Fe-S cluster assembly proteins CIAO1 and CIAO2B. **A.** Anti-His Western blot analysis of strep-tactin purified His-HBx either co-expressed with CIAO1 or expressed alone. **B.** Anti-His Western blot analysis of GSTrap purified CIAO2B co-expressed with both HBx and CIAO1 or CIAO1 alone. Note CIAO1 contains both N-terminal His- and Strep-tags, while CIAO2B carries both N-terminal His- and GST-tags. Separately purified CIAO1 and HBx (refolded) are also shown as controls. **C.** AlphaFold models of CIAO1 and the CIAO1:CIAO2B complexes in the absence (left) or presence (right) of HBx. Models are colored by pLDDT score, with HBx shown at 15% opacity.

To gain structural insight into the nature of these interactions, we used AlphaFold to model the CIAO1:HBx binary complex and the CIAO1:CIAO2B:HBx ternary complex (**Figs. 5C and S10**) (33). The models predict extensive electrostatic interactions between HBx and CIAO1. In the binary complex, HBx binds the narrow-end face of CIAO1’s β-propeller, a site recognized by client proteins of other WD-40 repeat proteins, and also CIAO2B (51–55). Upon inclusion of CIAO2B, this site is occupied, and HBx relocates to an adjacent region of CIAO1, spanning blades 3 and 4. Notably, blade 3 has been identified as a conserved binding site for other Fe-S client proteins (56). Similar interaction shifts were observed in AlphaFold models of other viral Fe-S cluster binding proteins, such as SARS-CoV-2 Nsp12 and Nsp13, further supporting the specificity of this binding mode (**Fig. S11**) (21, 22). Although HBx assumes a more compact conformation upon complex formation, the prediction confidence remained low (pLDDT < 50) for most HBx residues, regardless of the presence of CIAO1 alone or in complex with both CIAO1 and CIAO2B (**Figs. 1A, 5C, and S9C**). Importantly, in the predicted CIAO1:CIAO2B:HBx model, the key cysteine residues implicated in Fe-S cluster coordination (C61, C69, C143, and C148) are spatially proximal to, but not embedded within, the CIAO1/CIAO2B binding interface. This supports a model in which these cysteines primarily function in metallocofactor coordination, rather than mediating protein interactions within the CTC.

### Degradation of the HBx Fe-S cluster by nitroxide and NO donors

Given the essential role of HBx in maintaining HBV transcriptional activity, we next investigated whether its Fe-S cluster is susceptible to cell-permeable small molecules. Specifically, we tested the effect of the stable nitroxide TEMPOL and the NO donor DEA/NO, both of which are known to degrade Fe-S clusters in other proteins (22, 57, 58). Anaerobically purified DsbC-HBx with a [4Fe-4S] cluster was incubated with increasing amounts of TEMPOL or DEA/NO, and cluster integrity was assessed by EPR spectroscopy following chemical reduction with dithionite (**Fig. 6**). TEMPOL treatment led to a progressive loss of the [4Fe-4S]^+1^ cluster EPR signal, with ∼50% signal reduction observed at 24 eq and complete signal loss at 32 eq (per mol Fe per mol protein). No new EPR signals emerged following TEMPOL treatment, consistent with cluster disassembly and reduction of TEMPOL to its EPR-silent hydroxylamine form. In contrast, DEA/NO treatment induced a dose-dependent conversion of the Fe-S cluster into a distinct EPR-active species characteristic of dinitrosyl iron complexes (DNICs), a well-established degradation product of Fe-S clusters in the presence of NO(59). The DNIC signal was maximum at 2 eq of DEA/NO (corresponding to 3 eq of NO per mol Fe), indicating that NO-mediated disassembly was both more rapid and efficient than with TEMPOL (**Fig. 6**). A similar dose-dependent degradation pattern was observed for the [2Fe-2S] cluster form of DsbC-HBx following treatment with either TEMPOL or DEA/NO, demonstrating that both cluster types respond similarly to these nitroxides and NO donors (**Fig. S12**). The greater sensitivity to DEA/NO likely stems from the smaller size of the released NO when compared to TEMPOL, which likely allows easier access and more efficient degradation of the Fe-S cluster at lower concentrations. Together, these findings suggest that the Fe-S cluster in HBx is a redox-sensitive regulatory element and may represent a vulnerability that can be targeted to selectively disrupt HBx activity or modulate its function.

**Figure 6.**
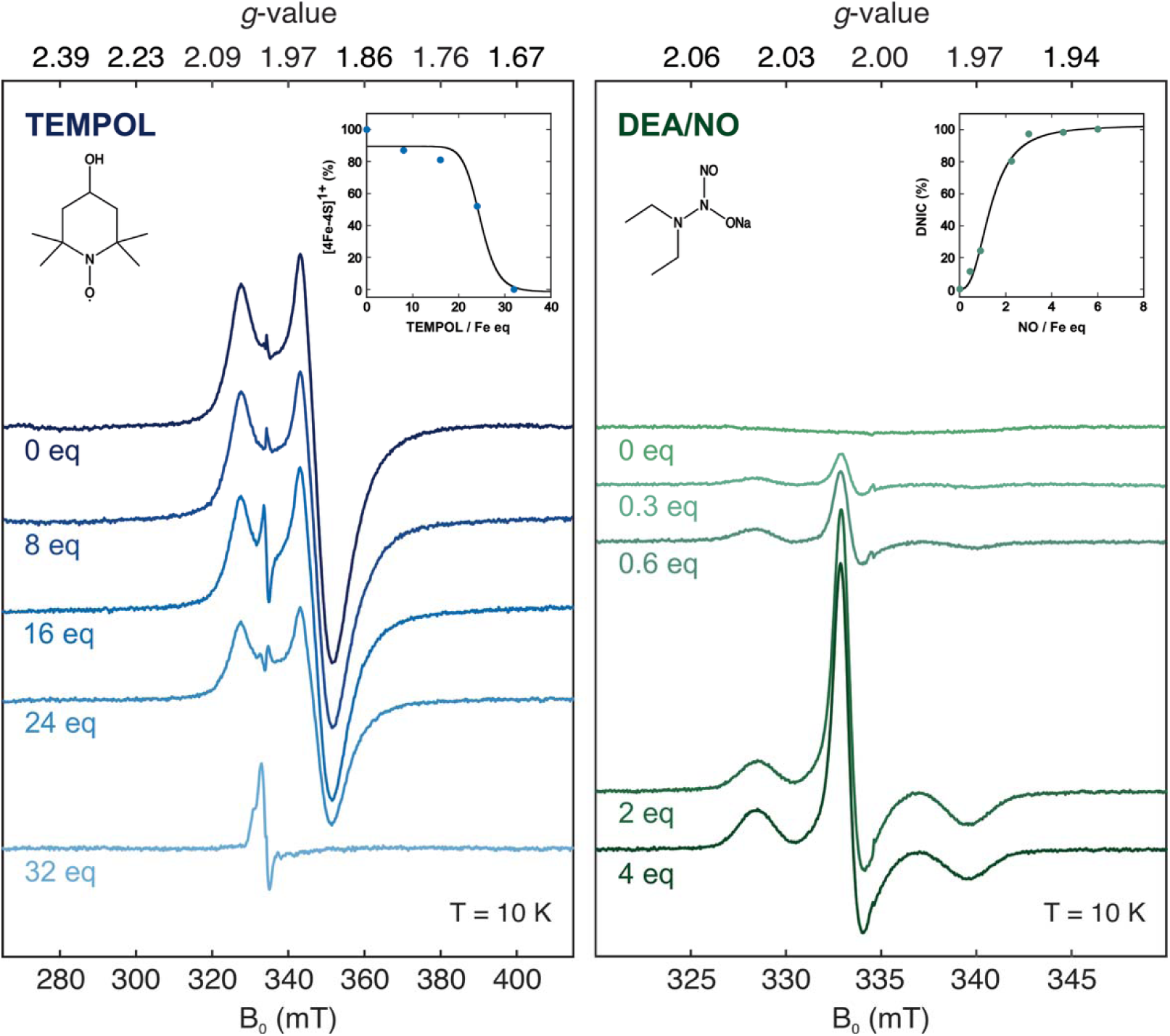
The nitroxide radical TEMPOL and the nitric oxide donor DEA/NO disassemble the Fe-S cluster of HBx. CW EPR spectra of [4Fe-4S]-cluster enriched DsbC-HBx obtained after incubation with increasing equivalents of TEMPOL (left) and DEA/NO (right, 1 mol DEA/NO liberates 1.5 mol NO). Spectra were recorded at a modulation amplitude = 1 mT and microwave power attenuation = 30 dB to monitor the TEMPOL-mediated disassembly of [4Fe-4S]^1+^ clusters, and 0.1 mT and 40 dB to monitor the NO-mediated DNIC formation. Additional experimental conditions: microwave frequency = 9.38 GHz, temperature = 10 K. Quantification was performed by double integration, and the integrated intensities of the [4Fe-4S]^1+^ cluster and DNIC signals were normalized to those at 0 eq TEMPOL and 4 eq DEA/NO, respectively. (Insets) The extent of [4Fe-4S]^1+^ cluster degradation or DNIC formation as a function of TEMPOL or NO equivalents with respect to the Fe content in HBx.

## Discussion

HBx was recently identified as an Fe-S cluster protein that coordinates a redox-active cofactor, capable of switching between an O_2_-stable [2Fe-2S] and an O_2_-sensitive [4Fe-4S] cluster form, with the latter proposed as the most likely physiologically relevant state *in cellulo* (6, 8). In addition to coordinating an Fe-S cluster, HBx can also bind Zn, as demonstrated by our study and others (6, 10). This dual metal binding ability raises a fundamental question about the chemical nature of the physiologically relevant metallocofactor *in vivo*. Here, we address this uncertainty and establish HBx as a *bona fide* Fe-S cluster protein, providing mechanistic insights into its metal-binding properties and how these can relate to its biological function.

We show that HBx interacts with the human cytosolic Fe-S cluster assembly machinery, specifically the targeting components CIAO1 and CIAO2B. These interactions provide compelling evidence that HBx engages the host biosynthetic machinery to acquire its cofactor, a feature that is essential given the fact that viruses lack the ability to synthesize such cofactors *de novo*. The identification of this interaction supports the physiological relevance of the Fe-S cluster in HBx and implies that disrupting its integrity could impair HBx function. This concept mirrors similar strategies applied to SARS-CoV-2, where targeting the Fe-S cluster in the RNA-dependent RNA polymerase Nsp12 inhibited viral replication (22). Similarly, we demonstrate that the Fe-S cluster in HBx is vulnerable to cell-permeable, low cytotoxicity redox-active small molecules. The nitric oxide donor DEA/NO fully disassembled both the [4Fe-4S] and [2Fe-2S] clusters at micromolar concentrations, while the stable nitroxide drug TEMPOL required higher (millimolar range) concentrations for complete degradation. Although these concentrations exceed reported IC_50_ values for DEA/NO and TEMPOL, this difference likely reflects limited small-molecule access to the Fe-S cluster *in vitro*, due to the use of a fusion-tagged HBx construct and its inherent oligomerization. Thus, the effective *in vivo* dose required for HBx-induced Fe-S cluster degradation may be lower (23, 60–63). Nevertheless, our findings expose a vulnerability of the HBx Fe-S cluster, regardless of its form, to both DEA/NO and TEMPOL, identifying them as promising therapeutic candidates for disrupting Fe-S cluster-dependent HBx function during HBV infection.

To enable selective targeting and rational drug design, it is critical to identify the ligands that are involved in the Fe-S cluster coordination. However, the lack of HBx sequence homology to known proteins, coupled with its high degree of structural disorder, makes inference of the binding site from computational structural models poorly reliable. By integrating mutagenesis studies with EPR spectroscopy, elemental analyses, and chemoproteomics, our study provides compelling evidence that HBx utilizes the conserved cysteines C61, C69, C143, and C148 as the most probable ligands for Fe-S cluster binding. C137 can also serve as a ligand, albeit with a lower probability, but this likelihood increases when some of these four cysteine ligands are replaced by alanines or serines. This behavior reflects the structural plasticity of the HBx polypeptide and aligns with our mutagenesis studies, in which neither single nor multiple cysteine substitutions were sufficient to completely abolish Fe-S cluster incorporation. This plurality of coordination modes can either be an actual functional property of HBx, enabling it to maintain Fe-S cluster binding, or an artifact resulting from our *in vitro* conditions and the absence of physiological interacting partners. Notably, the binding pattern remains consistent across both the [4Fe-4S] and [2Fe-2S] cluster forms, indicating the presence of a common binding site for these cofactors.

Our study further revealed that Fe-S clusters and Zn compete for a shared binding site in HBx, with C61, C69, C143, and C148 serving as primary ligands and C137 acting as a shared auxiliary ligand. Substitution of H139 with alanine had no effect on Fe incorporation, and HYSCORE spectroscopy also ruled out its involvement in Fe-S cluster coordination, indicating that Fe-S binding mainly relies on cysteines, potentially augmented by solvent-derived ligands. Elemental analyses were inconclusive regarding H139 being a Zn ligand, while chemoproteomics showed a cysteine ligation pattern similar to that for the Fe-S cluster. These experiments show that Zn ligation by H139 is less likely but cannot be ruled out. H139 may act as an additional Zn ligand, which could distinguish it from Fe-S cluster binding and imply functional differences influenced by cellular context and metal availability. Notably, C61, C69, C137, and H139 are each critical for HBx-mediated degradation of the Smc5/6 complex, pointing to a perhaps shared functional role for both metallocofactors in clinically relevant cysteine variants, but not in histidine variants (10, 30, 34). The observed diversity in both metallocofactor identity and coordination may provide HBx with a functional advantage, enhancing its metal-binding flexibility and supporting its activity under varying cellular conditions, thereby facilitating HBV’s chronic persistence in the host.

The ability of cysteine ligands to interchangeably coordinate Fe-S clusters and Zn is a common feature of many metalloproteins, supporting structural integrity and key cellular functions, such as electron transfer, redox balance, and metabolism (49, 64, 65). For HBx, mitochondrial targeting is likely independent of both the Fe-S cluster and Zn, as C115 is not involved in coordinating either cofactor (32). The common metal-binding ligands C61 and C69 are necessary for the interaction of HBx with DDB1, suggesting that both the Fe-S cluster and Zn may play a role in its transactivation function (30, 34). In addition, the shared auxiliary ligand C137 is vital for activating the signal transducer and activator of the transcription 3 (STAT3) pathway, implying that HBx bound to either metallocofactor may contribute to HCC progression during chronic HBV infection by upregulating this pathway (31, 66). On the other hand, our study demonstrated that mutations at the C-terminal cysteine residues C137, C143, and C148 significantly impair Fe-S cluster coordination. C-terminally truncated HBx variants, which are frequently found in chronic HBV patients (lacking 14-35 C-terminal residues, including C137, C143, and C148), exhibited a reduced ability to upregulate the hypoxia-inducible factor-1α (HIF-1α) (67). Moreover, the C-terminus is essential for HBx stability and transactivation and may also contribute to its role in hepatocellular carcinogenesis (35, 68). Overall, the Fe-S cluster ligands appear to be crucial for key aspects of HBx-mediated pathogenesis, underscoring the importance of the metallocofactor in HBx’s structure and function.

In summary, we define the likely ligands that coordinate both the Fe-S cluster and Zn in HBx, uncovering a fundamental but previously unrecognized connection between metallocofactor coordination and HBx function. Our findings support a model in which the Fe-S cluster serves as a structurally and functionally critical component of HBx, linking its metal-binding activity to host interactions and viral pathogenesis. The intrinsic disorder and metallocofactor binding observed in HBx may also be a generalizable feature of other viral proteins, particularly those that evade structural characterization. Thus, our approach provides a framework for identifying functional ligands in similarly flexible viral proteins. Although our conclusions are based on *in vitro* systems, they strongly suggest that the Fe-S cluster is a central regulator of HBx activity and represents a promising target for antiviral drug development.

## Materials and Methods

### Materials

All chemicals were of high-purity grade and obtained from Fisher Scientific unless otherwise specified.

### Generation of DsbC-HBx Variants

The HBx A2 sequence (WT or variant) was inserted into a pET-40b(+) vector (kindly gifted by Dr. Mehmet Berkmen, NEB) for its expression as a fusion with the disulfide bond isomerase (DsbC), which carried a kanamycin resistance marker. Single and multiple amino acid substitutions, as well as sequence truncations of HBx (genotype A2 with NCBI accession: P69713), were generated by Genscript USA Inc. (Piscataway, NJ) as fusions to an N-terminal DsbC to allow for soluble HBx.

### Expression of DsbC-HBx WT and Variants

The plasmids encoding DsbC-HBx (kanamycin resistance) were transformed into Rosetta (DE3) or T7 Express *Escherichia (E.) coli* competent cells (NEB, Ipswitch, MA) along with the pDB1282 plasmid that harbors genes involved in Fe-S cluster biosynthesis (ampicillin resistance, kindly gifted by Dr. Squire J. Booker, The Pennsylvania State University). Transformed cells were grown in Lennox Broth (LB) or minimal (M9) media at 37 °C with shaking (220 rpm) until the optical density at 600 nm (OD_600_) reached ∼0.3, at which time L-Arabinose (0.2% (w/v) final concentration) and L-cysteine (200 μM final concentration) were added. When cells reached an OD_600_ of ∼0.7, they were cold shocked at 4 °C for 1 hr. Protein expression was induced by the addition of 0.5 mM isopropyl β-d-1-thiogalactopyranoside (IPTG) and supplemented with 0.25 mM Fe(NH_4_)_2_(SO_4_)_2_ and/or ZnSO_4_ (no metal added for apoprotein expression). Cells were then incubated at 18 °C with shaking (220 rpm) for 18 to 20 hrs and harvested by centrifugation at 7,000 rpm for 15 min at 4 °C. Cell pellets were flash-frozen in liquid nitrogen (LN_2_) and stored at −80 °C prior to purification.

### Aerobic Isolation of DsbC-HBx WT and Variants

DsbC-HBx cell pellets were resuspended in Lysis Buffer A (50 mM HEPES, 300 mM NaCl, and 10 mM imidazole, 1 mM tris(2-carboxyethyl)phosphine (TCEP), pH 8.0). Phenylmethylsulfonyl fluoride (PMSF) was added to a final concentration of 45 μg/mL. Cells were lysed by sonication (QSonica) for 30 min (15 s pulse, 59 s pause, and 60% amplitude) and centrifuged at 22,000 x g for 30 min to remove cell debris. The clarified lysate was loaded onto a gravity-flow Ni-NTA column (McLab) equilibrated in Lysis Buffer A. The column was washed with Lysis Buffer A, followed by Wash Buffer A (50 mM HEPES, 300 mM NaCl, 30 mM imidazole, 1 mM TCEP, pH 8.0). Bound proteins were eluted with Elution Buffer A (50 mM HEPES, 150 mM NaCl, 300 mM imidazole, 10% glycerol, 1 mM TCEP, pH 8.0). Eluted fractions containing DsbC-HBx were pooled and concentrated using a 30 kDa MWCO Amicon Centrifugal Filter (Millipore, Sigma-Aldrich). The concentrated protein was further purified by size exclusion chromatography using a HiLoad 16/600 Superdex 200 pg column (GE Healthcare), which was equilibrated with Storage Buffer (50 mM HEPES, 150 mM NaCl, 10% glycerol, 1 mM TCEP, pH 8.0). Fractions containing the pure protein were combined and further concentrated. Protein purity was assessed via SDS-PAGE with Coomassie staining, and protein concentration was determined using the Bradford assay.

### Anaerobic Isolation of DsbC-HBx WT and Variants

DsbC-HBx was purified under strictly O_2_-free conditions in an anaerobic glovebox (Coy Labs, Grass Lake, MI) using degassed buffer solutions. Cell pellets were resuspended in Lysis Buffer A, followed by the addition of 45 μg/mL PMSF, 1 μg/mL DNase, 10 μg/mL lysozyme, 0.5 mM Fe(NH_4_)_2_(SO_4_)_2_, 5 mM 1,4-dithiothreitol (DTT), and 2 mM TCEP. This method allows for proper reconstitution of the [4Fe-4S] cluster in HBx (6). Cells were lysed by sonication for 30 min (15 s pulse, 59 s pause, 60% amplitude), during which time sodium sulfide (Na_2_S) was gradually added to a final concentration of 0.5 mM to promote Fe-S cluster formation. The centrifuged, clarified lysate was applied to a Ni-NTA column pre-equilibrated in Lysis Buffer A. Subsequent purification steps were performed as described above. The purified DsbC-HBx was stored in LN_2_ under anaerobic conditions until further use.

### Expression and Purification of the CIAO1:HBx Co-expressed Complex and HBx Alone

The plasmids encoding His_6_-tagged HBx (pET-28(+), kanamycin resistance) were either co-transformed with His_8_-TEV-Strep-tagged CIAO1 (pETDuet-1, ampicillin resistance) or transformed alone into T7 Express *E. coli* competent cells (69). Transformed cells were grown in LB medium at 37 °C with shaking (220 rpm) until an OD_600_ of ∼0.7 was reached. Cells were then cold shocked at 4 °C for 1 hr prior to induction with 0.5 mM IPTG. Following induction, cells were incubated at 18 °C for 18 to 20 hr, harvested by centrifugation, and stored at −80 °C until purification, as described above.

Cell pellets were resuspended in Lysis Buffer B (25 mM 2-(N-morpholino)ethanesulfonic acid (MES), 0.1% Nonidet P-40 (NP-40), pH 7.0) supplemented with 45 μg/mL PMSF, 1 μg/mL DNase, and 10 μg/mL lysozyme. Cells were lysed using a microfluidizer (LM20, Microfluidics), and the clarified lysate was applied to a strep-tactin column (IBA Lifesciences) equilibrated with Lysis Buffer B. The column was washed with Lysis Buffer B, followed by Wash Buffer B (25 mM MES, pH 7.0), and bound proteins were eluted using Elution Buffer B (25 mM MES, 2.5 mM *d*-Desthiobiotin, pH 7.0). Eluted fractions were analyzed by SDS-PAGE with Coomassie staining, and protein-containing fractions were concentrated using the 10 kDa MWCO Amicon Centrifugal Filter (Millipore, Sigma-Aldrich).

### Expression and Purification of the CIAO1:CIAO2B:HBx and CIAO1:CIAO2B Co-expressed Complexes

The plasmids encoding both His_6_-GST-TEV-tagged CIAO2B and His_8_-TEV-Strep-tagged CIAO1 (pETDuet-1, ampicillin resistance) were either co-transformed with His_6_-tagged HBx or transformed alone into T7 Express *E. coli* competent cells. Transformed cells were expressed, harvested, and stored for purification as described for the CIAO1:HBx co-expressed complex. Cell pellets were resuspended and lysed as described for the CIAO1:HBx co-expressed complex. The clarified lysate was applied to a GSTrap™ 4B column (Cytiva) equilibrated with Lysis Buffer B. The column was washed with Lysis Buffer B, and bound proteins were eluted using Elution Buffer C (25 mM MES, 10 mM L-glutathione (reduced), pH 7.0). Eluted fractions were analyzed and concentrated as described for the CIAO1:HBx co-expressed complex.

### EPR Spectroscopy

All EPR samples were prepared in Storage Buffer under O_2_-free conditions in an anaerobic glovebox. Samples were reduced with 6 mM sodium dithionite for 20 min at room temperature prior to being frozen in LN_2_, unless stated otherwise.

X-band continuous-wave (CW) EPR measurements were performed on a Bruker EleXsys E500 EPR spectrometer (operating at approximately 9.38 GHz) equipped with a rectangular resonator (TE_102_) and a continuous-flow cryostat (Oxford 910) with a temperature controller (Oxford ITC 503). The CW EPR spectra were recorded at 10 K, with a microwave power of 0.2 mW and a modulation amplitude of 1 mT, unless stated otherwise.

Pulse EPR measurements were performed on a Bruker Elexsys E580 X-band spectrometer equipped with a SuperX-FT microwave bridge. A Bruker ER4118X-MS5 resonator was used in combination with an Oxford CF935 helium flow cryostat. Microwave pulses generated by the microwave bridge were amplified by a 1 kW traveling wave tube (TWT) amplifier (Applied Systems Engineering, model 117x). The HYSCORE spectra were recorded at 10 K, with a magnetic field of 345 mT, microwave frequency of 9.44 GHz, *τ* = 132 ns, and *t*(π/2) = 8 ns.

### Metal Quantification

Metal analysis of the purified proteins was performed using inductively coupled plasma atomic emission spectrometry (ICP-AES) at the Laboratory for Isotopes and Metals in the Environment (LIME) at Pennsylvania State University (University Park, PA). For the sample preparation, 5 μM of protein was mixed with 2.5 mL of 7% nitric acid (HNO_3_, trace metal grade) solution and water to a final volume of 5 mL in a 15 mL conical tube. The sample solution was incubated at room temperature for 24 hrs to allow complete protein precipitation. After incubation, samples were centrifuged and filtered to remove precipitates and particulates. The resulting clear sample solution was sent for ICP-AES analysis.

### Fe-S cluster and Zn Reconstitution

All reconstituted Fe-S cluster and Zn samples were prepared in an anaerobic glovebox using anaerobically as-purified DsbC-HBx protein that was isolated from cells expressed in M9 media in the absence of any exogenously supplemented metal ions to obtain the apo form with degassed Storage Buffer. All reagent concentrations reported below are relative to the determined protein concentration.

For Fe-S cluster reconstitution, the protein samples were reduced with a 10-fold excess of DTT in the Storage Buffer and incubated at room temperature for 30 min. The *E. coli* IscS was expressed as a fusion with an N-terminal SUMO tag (the plasmid of pSUMO-IscS encodes for kanamycin resistance and was kindly gifted by Dr. Squire J. Booker) and isolated as described previously (70). A 50-fold excess of PLP-loaded IscS was added to the protein samples. For holo-Fe-S cluster samples, four molar equivalents of FeCl_3_ were added slowly over the course of 1 hr, followed by the addition of L-cysteine. No Fe and cysteine were added for the apo-Fe-S cluster samples. Both holo and apo samples were incubated overnight at 4 °C. The reconstitution mixtures were spun to remove any precipitate, and the supernatant was applied to a PD-10 desalting column to eliminate adventitious iron prior to isotopic N-ethylmaleimide (NEM) labeling.

For Zn reconstitution, the protein samples were reacted with one molar equivalent of ZnSO_4_ (holo-Zn) or an equal volume of water (apo-Zn) in the Storage Buffer. The samples were incubated at room temperature for 1 hr prior to isotopic NEM labeling.

### Isotopic NEM-labeling and MS Sample Preparation

In an anaerobic glovebox, 100 µL of apo (1mg/mL) and [4Fe-4S] or Zn reconstituted holo-HBx (1mg/mL) was labeled with 1 mM (1 µL of 100 mM stock) of either isotopically light NEM-d_0_ (apo-HBx) or heavy NEM-d_5_ (holo-HBx), mixed, and incubated at room temperature for 1 hr. The protein samples were frozen and stored at −80 °C until further processing. This procedure was repeated for [2Fe-2S] HBx under aerobic conditions.

Paired protein samples of NEM-labeled apo- and holo-HBx were thawed on ice and immediately precipitated with 5% TCA (5 µL of 100% TCA in water). After the addition of TCA, samples were vortexed and stored at −80 °C for at least 2 hrs or overnight. Samples were then thawed on ice and centrifuged at 15,000 x g for 10 min at 4 °C to collect precipitated protein. The supernatant was removed, and the protein pellet was resuspended in 500 µL of ice-cold acetone by vortexing and sonication. The protein precipitate was pelleted by centrifugation at 5,000 x g for 10 min at 4 °C. The supernatant was removed, and the protein pellet was allowed to air-dry until the acetone had completely evaporated. The protein precipitate was resolubilized in 30 µL of urea (8 M fresh stock in PBS) by sonication, followed by the addition of 70 µL of ammonium bicarbonate (100 mM fresh stock in water). To reduce any protein thiols, 1.5 µL of DTT (150 mg/ml fresh stock in water) was added, and samples were incubated at 75 °C for 15 min. To alkylate free thiols, 2.5 µL of iodoacetamide (94 mg/ml fresh stock in water) was added to samples, followed by incubation at room temperature for 30 min in the dark. The protein samples were then diluted with 120 µL of PBS. To digest protein samples, 2.5 µL of calcium chloride (100 mM stock in water) and 4 µL of sequencing grade trypsin (resuspended at 0.5 µg/µl in the provided trypsin resuspension buffer) (Promega) were added, and the samples were incubated overnight at 37 °C. The samples were acidified by the addition of 12.5 µL of MS-grade formic acid and combined pairwise (light NEM-d_0_ labeled apo-HBx with heavy NEM-d_5_ labeled holo-HBx) into a clean Eppendorf tube (final volume ∼ 500 µL). Samples were desalted using Sep-Pak C18 cartridges, which had been conditioned with 3 x 1 ml 100% acetonitrile and equilibrated with 3 x 1 ml Buffer A (5% acetonitrile, 0.1% formic acid). Samples were loaded onto the cartridges and allowed to flow through under ambient pressure. Samples were then reloaded onto the cartridge to improve yield. Cartridges were washed with 3 x 1 ml Buffer A. Samples were eluted from the cartridge into LoBind Eppendorf tubes using 2 x 0.5 ml Buffer B (80% acetonitrile, 0.1% formic acid). Samples were evaporated to dryness in a SpeedVac and stored at −20 °C until LC-MS/MS analysis. Prior to analysis, samples were resuspended in 500 µL of Buffer A.

### LC-MS/MS Analysis

Samples were analyzed by LC-MS/MS using an Exploris 240 mass spectrometer (Thermo Scientific) coupled to a Dionex Ultimate 3000 RSLCnano system. Samples (5 µL) were directly injected onto an Acclaim PepMap 100 (Thermo Scientific) loading column. Peptides were eluted onto an Acclaim PepMap RSLC (Thermo Scientific) and separated with a 1 hr gradient of Buffer B (20% water, 80 % acetonitrile, 0.1% formic acid) in Buffer C (100% water, 0.1% formic acid) at a flow rate of 0.3 µL/min. The spray voltage was set to 2.1 kV. One full MS1 scan (120,000 resolution, 350-1800 m/z, RF lens 65%, AGC target 300%, automatic maximum injection time, profile mode) was obtained every 2 s with dynamic exclusion (repeat count 2, duration 10 s), isotopic exclusion (assigned), and apex detection (30% desired apex window) enabled. A variable number of MS2 scans (15,000 resolution, AGC 75%, maximum injection time 100 ms, centroid mode) were obtained between each MS1 scan based on the highest intensity precursor masses, filtered for monoisotopic peak determination, theoretical precursor isotopic envelope fit, intensity (5E4), and charge state (2–6). MS2 analysis consisted of the isolation of precursor ions (isolation window 2 m/z) followed by higher-energy collision dissociation (HCD, collision energy 30%). Each sample was subjected to a second injection to generate two technical replicates per biological sample. All LC-MS/MS data were collected for multiple (2–5) biological replicates.

### Mass Spectrometry Data Processing

The tandem MS data were analyzed by the Thermo Proteome Discoverer V2.4 software package and searched using the SequestHT and Percolator algorithms against a UniProtKB database (www.uniprot.org) of the *E. coli* K12 proteome supplemented with the WT HBx protein sequence (genotype A2). Trypsin was specified as the protease with a maximum of 2 missed cleavages. Peptide precursor mass tolerance was set to 10 ppm with a fragment mass tolerance of 0.02 Da. Oxidation of methionine (+15.995) and modification of cysteine by alkylation (+57.021) or either isotopically light (+125.048) or heavy (+130.079) NEM were set as dynamic modifications, while acetylation (+42.011) and/or methionine-loss (+131.040) of the protein N-terminus were set as static modifications. The false discovery rate (FDR) for peptide identification was set to 1%. Light/heavy (L/H) ratios for cysteine-containing HBx peptides were calculated as the ratio of the NEM-d_0_ over NEM-d_5_ modified peptide precursor ion intensities. Quantified peptides were filtered out from the final dataset if they did not meet the following criteria: (1) have a minimum of 3 PSMs, (2) have good coverage across replicates and consist of majority non-singleton values (e.g., ion intensity was observed for both the light and heavy peptide species across most replicates), (3) have all cysteine residues accurately and exclusively assigned as NEM-labeled, and (4) have a minimal number of missed tryptic cleavages, especially where these would introduce additional cysteine residues to a peptide. L/H ratios were then culled of remaining singleton values, and each L/H ratio was corrected by the associated median L/H ratio for that replicate, generating a final high-confidence dataset of corrected L/H ratios for each HBx cysteine-containing peptide.

### Western Blotting

For anti-His detection, membranes were probed with a 1:2000 dilution of alkaline phosphatase conjugated monoclonal mouse anti-6X His antibody (Abcam) and developed using 1-Step™ NBT/BCIP Substrate Solution. For anti-Strep II detection, membranes were probed with a 1:5000 dilution of horseradish peroxidase (HRP) conjugated monoclonal anti-Strep II antibody (Millipore, Sigma-Aldrich) and developed using 1-Step™ Ultra TMB-Blotting Solution. For anti-GST detection, membranes were probed with a 1:5000 dilution of HRP conjugated monoclonal anti-GST antibody (GenScript) and developed using 1-Step™ Ultra TMB-Blotting Solution. All blots were visualized using a ChemiDoc MP Imaging System (Bio-Rad).

### Mass Spectrometry of CIAO1, CIAO2B, and HBx Proteins

Gel bands were excised and stored in Eppendorf tubes containing 500 µL of 100 mM ammonium bicarbonate at 20□°C until processing. The gel bands were washed for 15 min in 100 mM ammonium bicarbonate, then incubated in 200 µL of 10 mM TCEP at 60 °C for 30 min. After discarding the supernatant, the gel bands were incubated with 200 µL of 55 mM iodoacetamide in the dark at room temperature for 30 min. After discarding the supernatant, the gel bands were washed 3 times in 500 µL of 100 mM ammonium bicarbonate in 50:50 water:acetonitrile, once with 50 µL of acetonitrile, and dried in a SpeedVac for 5 min. In-gel digestion was performed by the addition of 20 µL of 10 ng/µL trypsin in 25 mM ammonium bicarbonate (200 ng trypsin total) for 10 min to rehydrate the gel slices, followed by the addition of enough 25 mM ammonium bicarbonate to cover the gel slice. The in-gel digests were incubated overnight at 37 °C, after which the supernatant was transferred to a clean tube, and the gel slices were incubated with 5% formic acid for 15 min at room temperature. The supernatant was removed and combined in the same tube as the in-gel digestion supernatant. The gel slices were then washed 3 times with 100 µL of acetonitrile for 15 min, and each time the supernatant was added to the collection tube with the initial in-gel digestion supernatant. Samples were evaporated to dryness in a SpeedVac and stored at −20 °C until LC-MS/MS analysis. Prior to analysis, samples were resuspended in Buffer A and analyzed by LC-MS/MS as described above. The tandem MS data were analyzed as previously described with the following modifications: (1) human recombinant CIAO1 and CIAO2B protein sequences were added to the protein search database, (2) only oxidation of methionine (+15.995) and modification of cysteine by alkylation (+57.021) were applied as dynamic modifications, and (3) identified proteins with less than 2 peptides and fewer than 20 PSMs were filtered out from the results.

### Fe-S Cluster Degradation Assay

The assays were conducted inside an anaerobic glovebox using degassed buffer solutions. The TEMPOL (4-hydroxy-2,2,6,6-tetramethylpiperidin-1-oxyl) was dissolved in Storage Buffer to prepare a 25 mM stock solution. The DEA/NO (Diethylammonium (Z)-1-(N,N-diethylamino)diazen-1-ium-1,2-diolate) was initially dissolved in 50 mM NaOH solution to make a 100 mM solution, then diluted with Storage Buffer to generate a 10 mM stock solution. The DEA/NO solutions were set at room temperature for 1.5 hrs (*t_1/2_* = 16 min at 22-25 °C) to allow NO release to reach saturation prior to protein incubation. Various equivalents of TEMPOL or DEA/NO were incubated with anaerobically or aerobically isolated DsbC-HBx proteins containing either [4Fe-4S] or [2Fe-2S] clusters, respectively, for 15 min as indicated in the main and supplementary text and figure legends. Samples were prepared for EPR spectroscopy as described above.

## Data Availability

All data supporting the findings of this study are available within the article and its Supplementary Information files. Any additional data is available from the corresponding author upon reasonable request.

## Acknowledgments

The authors thank Dr. Paul Ralifo for providing us access to the EPR spectrometer at the Chemical Instrumentation Center of Boston University (Boston, MA). The authors thank Dr. Laura J. Liermann for ICP-AES analysis at the Laboratory for Isotopes and Metals in the Environment at Pennsylvania State University (University Park, PA). The authors are also thankful to Dr. Mehmet Berkhmen (New England Biolabs) for donating the backbone of the DsbC vector. The authors thank Dr. Bruce Goode (Brandeis University) for providing unrestricted access to their microfluidizer. The authors would also like to thank Dr. Julia Kardon (Brandeis University) for providing the Western blot equipment and her graduate student, Cori Posner, for assisting us in troubleshooting the Western blot experiments. The authors are additionally grateful to Amy Milne for her contributions and helpful discussions.

