## supporting Information for "Promiscuous Metal Site in Hepatitis B Virus X Protein Binds an Fe-S Cluster"

Jiahua Chen^1^, Michelle Langton^1^, Patrick Cao^1^, Avital Aaron^1^, Jackson Ho^2^, Eranthie Weerapana^3^, Deborah L. Perlstein^2^, Alexey Silakov^4^, Daniel W. Bak^3*^, Maria-Eirini Pandelia^1^*

^1^ Department of Biochemistry, Brandeis University, Waltham, Massachusetts 02453, United States

^2^ Department of Chemistry, Boston University, Boston, Massachusetts 02215, United States

^3^ Department of Chemistry, Boston College, Chestnut Hill, Massachusetts 02467, United States

^4^ Department of Chemistry, The Pennsylvania State University, University Park, Pennsylvania 16802, United States

*Corresponding authors

**This PDF file includes:**

Figures S1 to S12

Tables S1 to S4

SI Reference

**Figures**

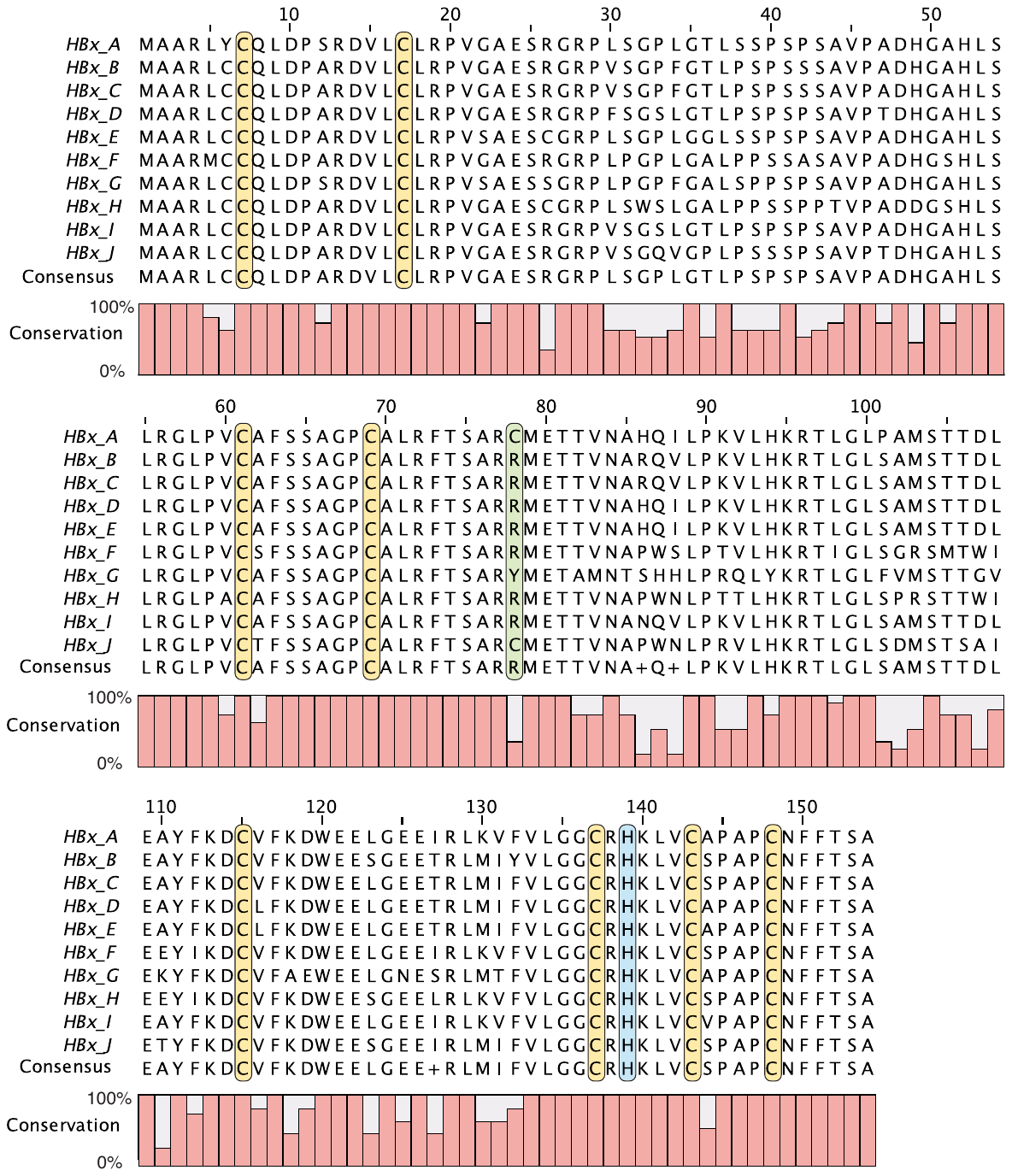

**Fig. S1.** **Multiple sequence alignment of HBx from the 10 HBV genotypes A to J.** Cysteine and histidine residues conserved across all sequences are highlighted in yellow and blue, respectively. The residue at position 78 is highlighted in green, showing a cysteine present only in HBV genotypes A and J, which is a tyrosine in genotype G and an arginine in genotypes B-F and H-I. The consensus sequence of HBx and the residue conservation bar plot are also shown.

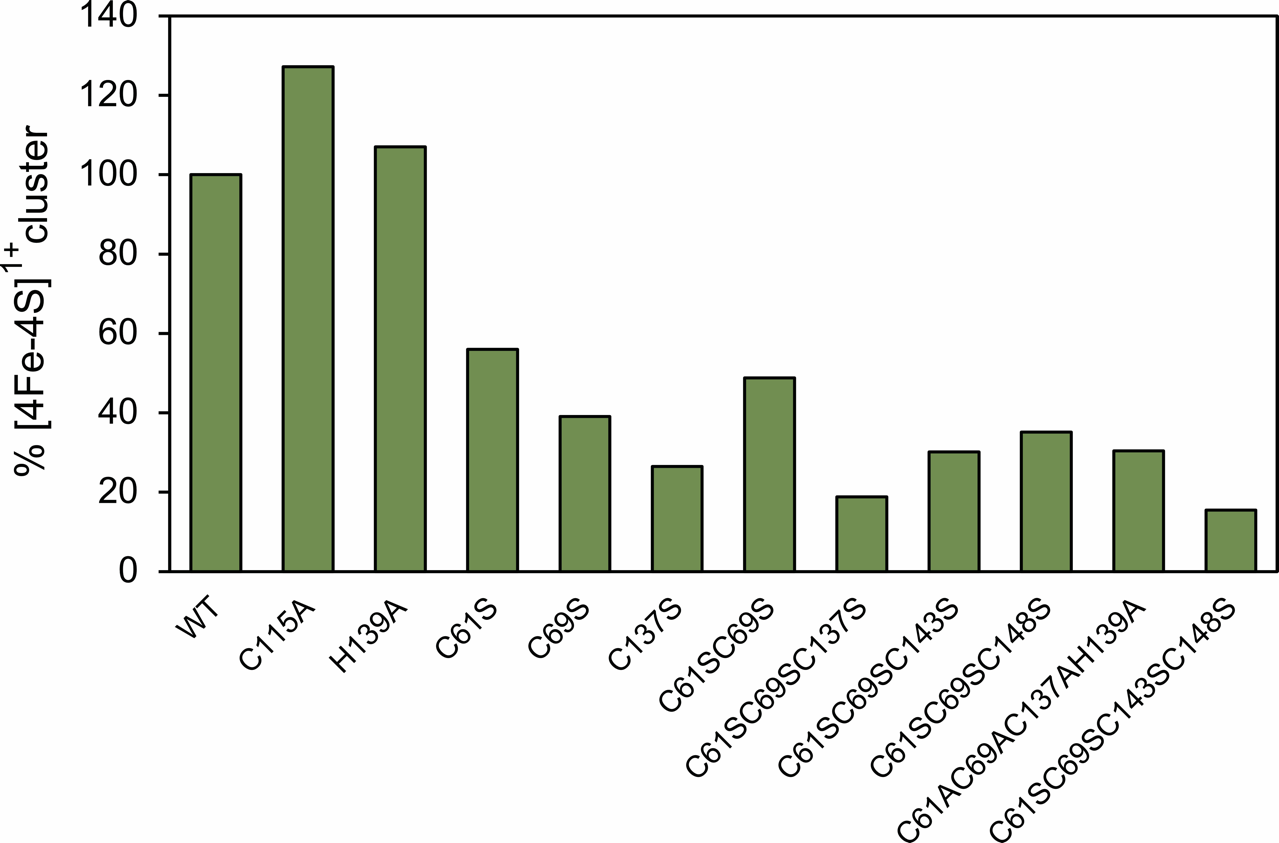

**Fig. S2. Extent of [4Fe-4S]^1+^ cluster incorporation in WT and variant DsbC-HBx.** The amount of [4Fe-4S]^1+^ cluster present was quantified by double integration of the first-derivative EPR spectra, with signal intensities normalized to account for the same protein concentration. The integrated area of the WT [4Fe-4S]^1+^ cluster signal was considered 100%, and the areas of the variant spectra were normalized with respect to that of the WT protein. The bar graph is a quantitative representation of the extent of Fe-S cluster incorporation in the variant DsbC-HBx with respect to that of the WT protein, as shown in **Figure 1** of the main manuscript.

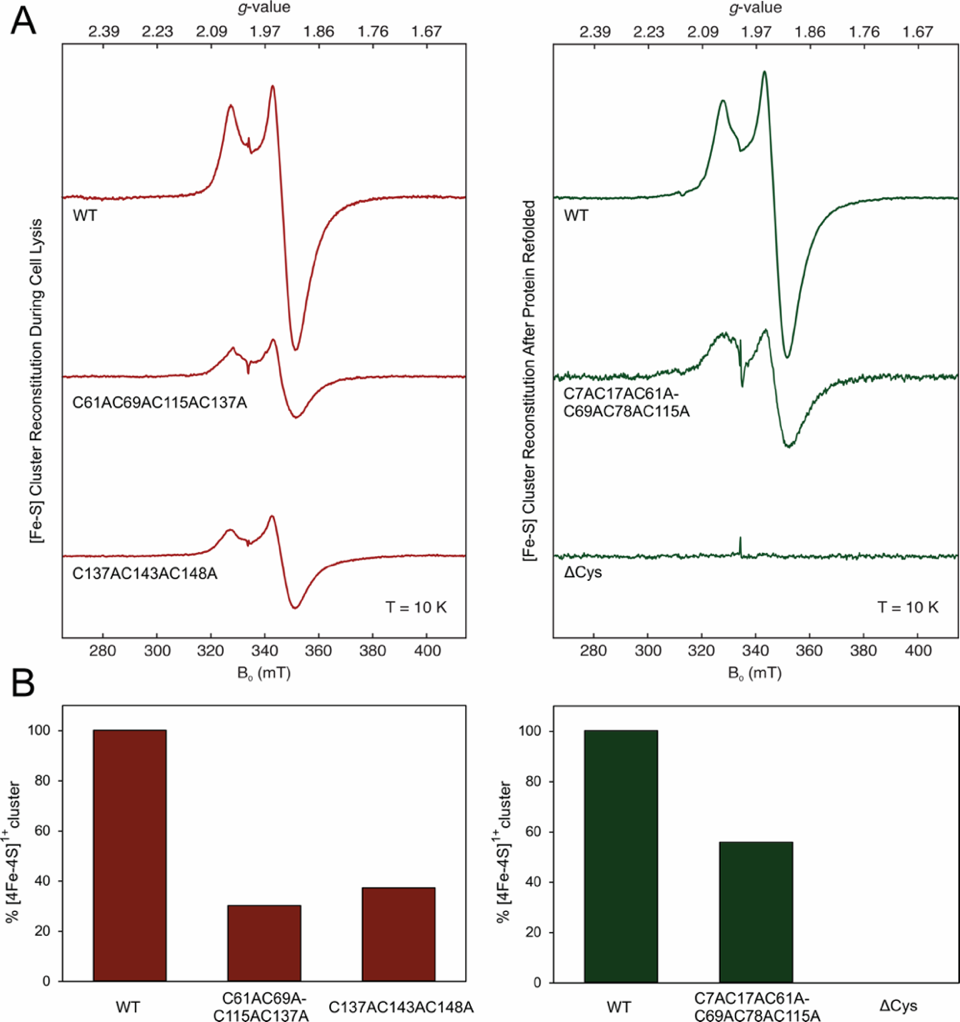

**Fig. S3.** **X-band** **CW EPR spectra of the WT and variant DsbC-HBx monitoring [4Fe-4S]^1+^ cluster assembly.** WT and variant DsbC-HBx were co-expressed with the pDB1282 plasmid in M9 media supplemented with Fe(NH_4_)_2_(SO_4_)_2_, then isolated under native conditions (left) or refolded following isolation under denaturing conditions (right) and was either chemically reconstituted during lysis or protein refolding, respectively (protein refolding method described in SI ref. 1). Irrespective of the isolation/reconstitution method, both approaches yielded comparable enrichment of a [4Fe-4S] cluster. This validation step was necessary because some variants were insoluble and needed to be reconstituted after refolding to assess their ability to incorporate an Fe-S cluster. Multiple-point N-terminal (C7AC17AC61AC69AC78AC115A - insoluble) and C-terminal (C61AC69AC115AC137A and C137AC143AC148A - soluble) variants all assembled a [4Fe-4S] cluster with varying efficiencies. Only substitution of all cysteines with alanines (ΔCys variant - insoluble) resulted in a complete loss of the protein’s ability to coordinate an Fe-S cluster. **A.** CW EPR spectra of WT and variant DsbC-HBx. The intensities have been normalized to account for the same protein concentration. Experimental conditions: microwave frequency = 9.38 GHz, temperature = 10 K, microwave power = 0.64 mW, modulation amplitude = 1 mT. **B.** Bar graph representing the extent of [4Fe-4S] cluster incorporation in WT and variant DsbC-HBx after double integration of the first-derivative EPR signals. The area corresponding to the WT [4Fe-4S]^1+^ cluster signal was considered 100%, and the areas of the variant EPR spectra were normalized with respect to that of the WT DsbC-HBx.

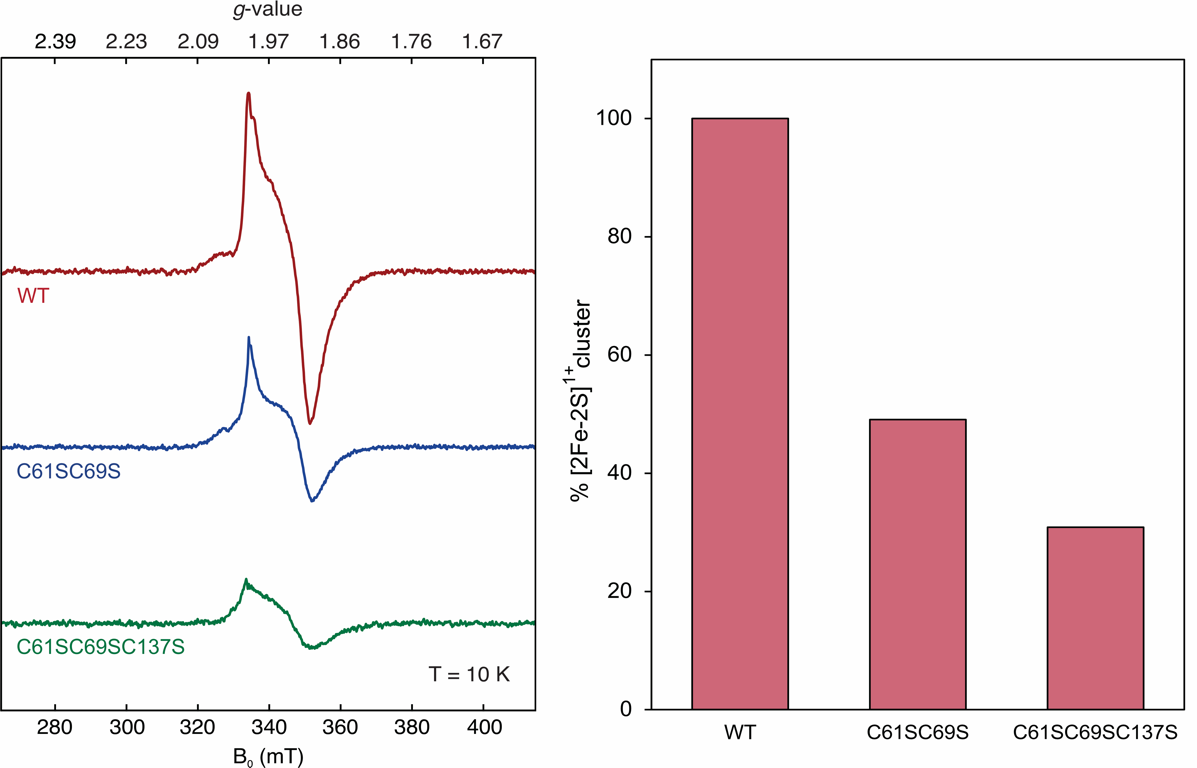

**Fig. S4.** **X-band** **CW EPR spectra of WT and variant DsbC-HBx monitoring [2Fe-2S]^1+^ cluster assembly**. WT and variant DsbC-HBx were co-expressed with the pDB1282 plasmid in M9 media supplemented with Fe(NH_4_)_2_(SO_4_)_2_, then aerobically isolated with a [2Fe-2S] cluster. The transient one-electron reduced [2Fe-2S]^1+^ form was trapped by reduction with excess sodium dithionite (6mM), followed by rapid freezing after 30 s. (Left) EPR spectra of WT and variant DsbC-HBx. The EPR spectral intensities have been normalized to account for the same protein concentration. Experimental conditions: microwave frequency = 9.38 GHz, temperature = 10 K, microwave power = 0.64 mW, modulation amplitude = 1 mT. (Right) The percentage of [2Fe-2S]^1+^ cluster was estimated after double integration of the first derivative EPR spectra, with signal intensities normalized to account for the same protein concentration. The cluster percentage is normalized with respect to that of the WT, which is considered 100%.

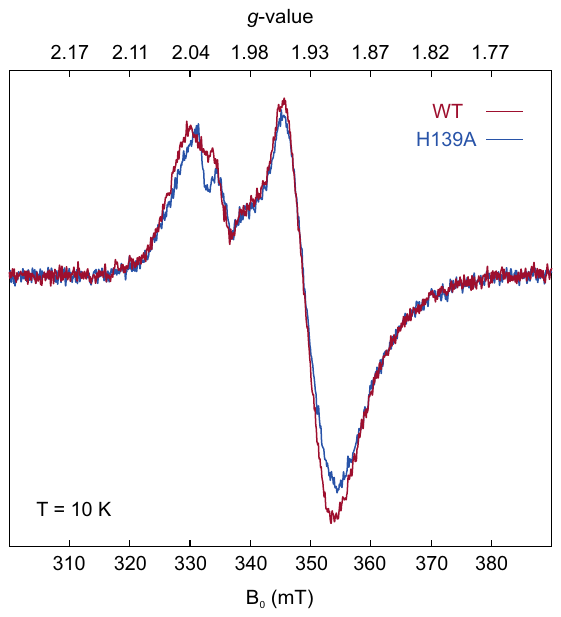

**Fig. S5. X-Band CW EPR spectra of WT and H139A DsbC-HBx.** WT and H139A DsbC-HBx were co-expressed with the pDB1282 plasmid in M9 media supplemented with Fe(NH_4_)_2_(SO_4_)_2_, then isolated under O_2_-free conditions and chemically reconstituted during lysis to obtain the [4Fe-4S]^2+^ cluster. The purified proteins were reduced with an excess of sodium dithionite (6 mM) for 20 min, which exhibit EPR spectra characteristic of a [4Fe-4S]^1+^ cluster, with principal values of *g*_II_ = 2.04 and *g*_⊥_=1.94. Experimental conditions: microwave frequency = 9.44 GHz, temperature = 10 K, microwave power = 1 mW, modulation amplitude = 1 mT.

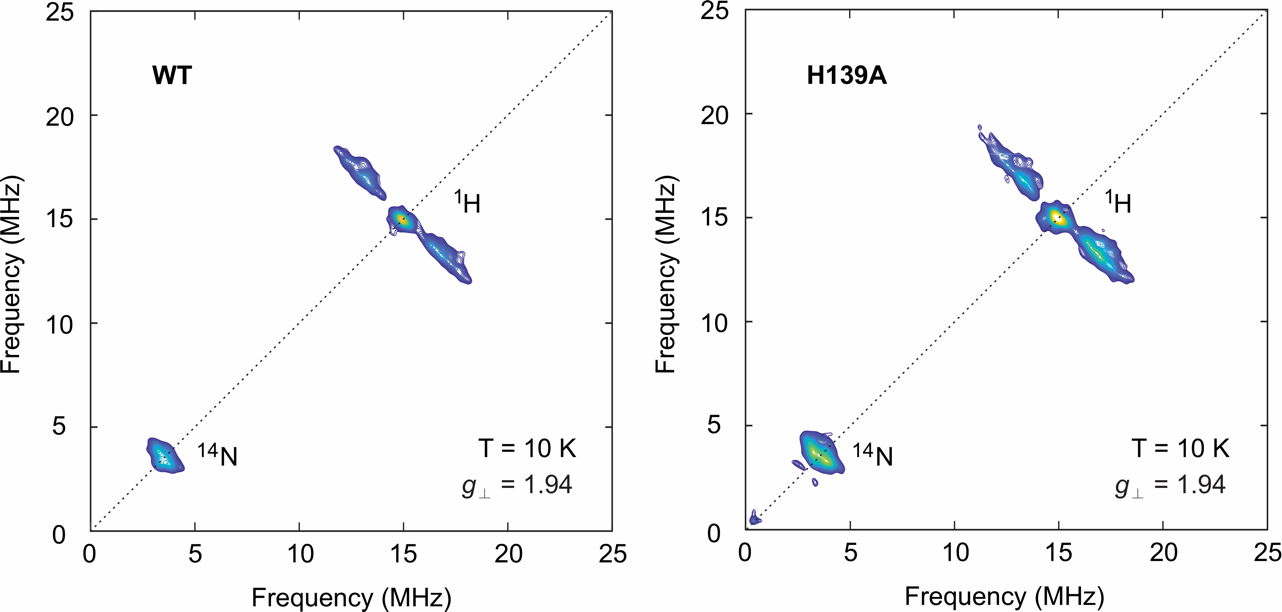

**Fig. S6. X-band HYSCORE spectra of cryo-reduced WT and H139A DsbC-HBx**. WT (left) and H139A (right) DsbC-HBx were co-expressed with the pDB1282 plasmid in M9 media supplemented with Fe(NH_4_)_2_(SO_4_)_2_, then isolated under O_2_-free conditions and chemically reconstituted during lysis to obtain the [4Fe-4S]^2+^ cluster. The paramagnetic [4Fe-4S]^1+^ form was generated by cryo-reduction using a Co^60^ source at liquid nitrogen temperatures. The HYSCORE spectra were recorded at a magnetic field of 345 mT, corresponding to the principal value *g*_⊥_ of 1.94. Experimental conditions: microwave frequency = 9.44 GHz, temperature 10 K, *τ* = 132 ns, *t*(π/2) = 8 ns.

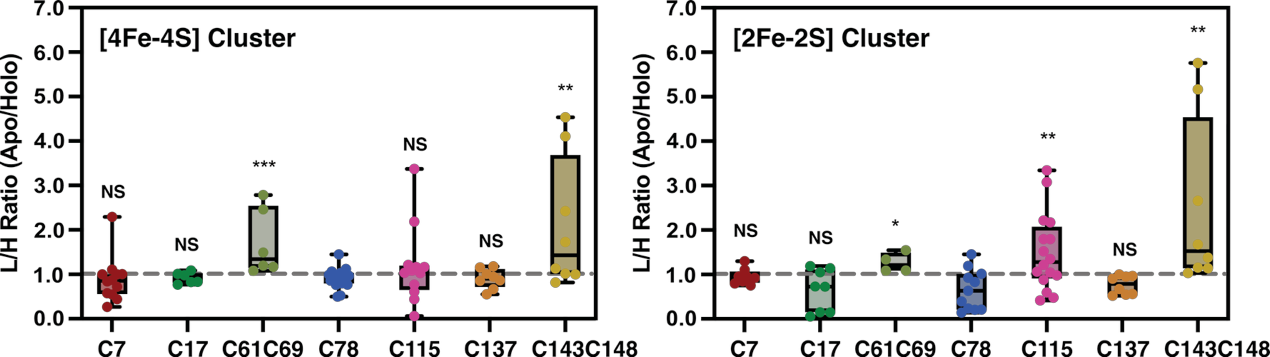

**Fig. S7. Chemoproteomics shows the same Fe-S cluster-binding cysteines in HBx for both [4Fe-4S] and [2Fe-2S] cluster forms.** DsbC-HBx was expressed in the absence (apo) or the presence (holo) of exogenously added Fe(NH_4_)_2_(SO_4_)_2_, followed by purification and isotopic labeling with the cysteine-specific alkylating reagent, NEM. The isotopically labeled apo- and holo-HBx samples were subjected to TCA precipitation, DTT reduction, alkylation, and trypsin digestion, then combined pairwise (apo and holo) for LC-MS/MS analysis to obtain the cysteine reactivity expressed as L/H ratios. The L/H ratios for independently expressed and isolated [4Fe-4S] (left) and [2Fe-2S] (right) cluster-containing proteins are shown as box plots. For each cysteine residue, each dot represents an independent L/H ratio measurement. The median L/H ratio value across all cysteine residues (C7, C17, C61C69, C78, C115, C137, and C143C148) is indicated by a dashed line. Statistical significance was determined by an independent *t*-test and is denoted as NS (not significant), **p* < 0.05, ***p* < 0.01, and ****p* < 0.001.

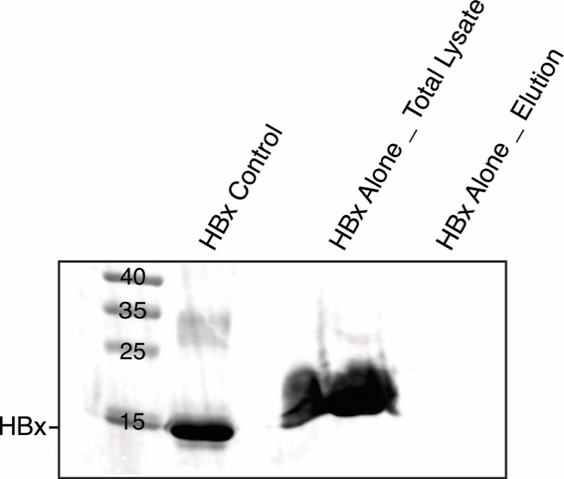

**Fig. S8. HBx is insoluble when expressed without a solubility enhancement tag.** Anti-His Western blot analysis of His-HBx expressed without a solubility tag (i.e., DsbC) and purified via Ni-NTA chromatography under native conditions. His-HBx obtained via a denaturing purification step, and refolding is shown as a control (protein refolding method described in SI ref. 1).

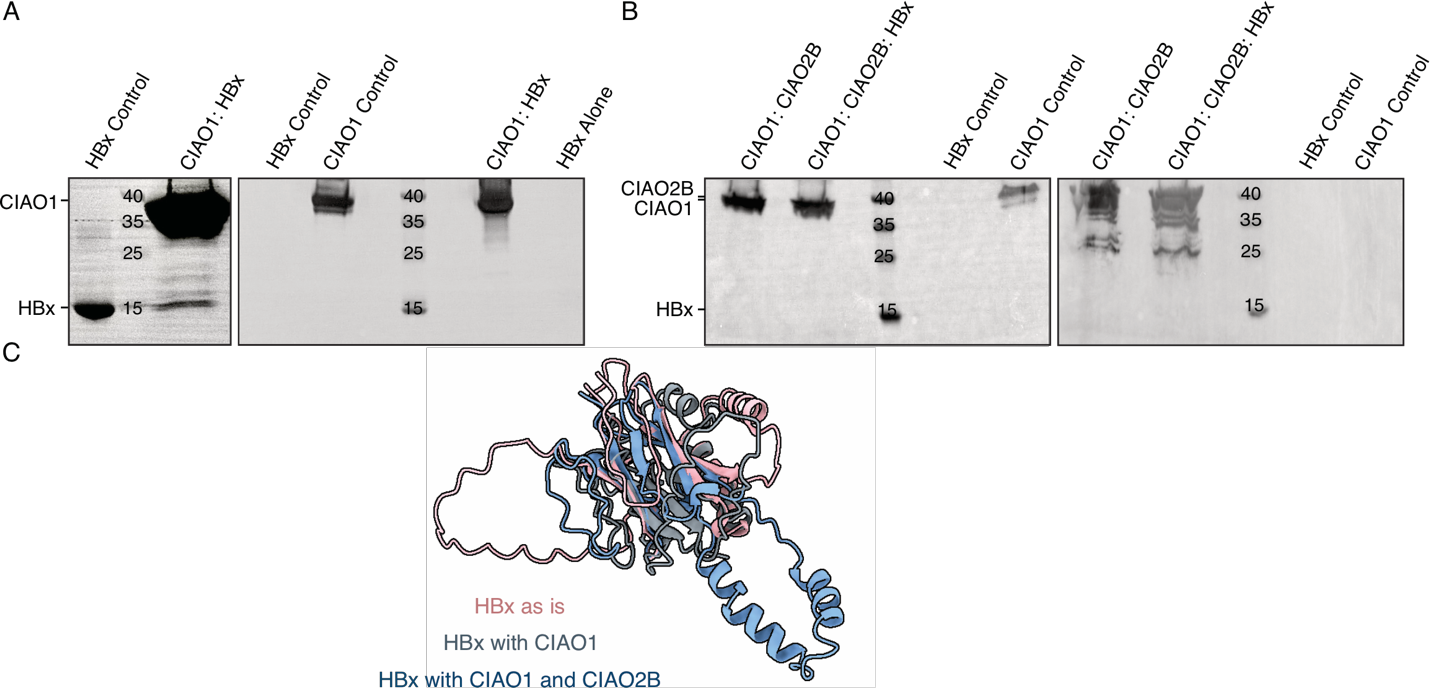

**Fig. S9. HBx interacts with human cytosolic Fe-S cluster assembly proteins CIAO1 and CIAO2B. A.** Anti-His (left) and anti-Strep (right) Western blot analysis of aerobic strep-tactin purified His-HBx co-expressed with CIAO1 or expressed alone. **B.** Anti-Strep (left) and anti-GST (right) Western blot analysis of aerobic GSTrap purified CIAO2B co-expressed with both His-HBx and CIAO1 or with CIAO1 alone. Separately purified CIAO1 and His-HBx (refolded) are also shown as controls. **C.** Superimposed AlphaFold models of HBx as is (pink), in complex with CIAO1 (grey), and in complex with both CIAO1 and CIAO2B (blue) to probe any predicted structural changes imposed by the presence of one or both CIA proteins.

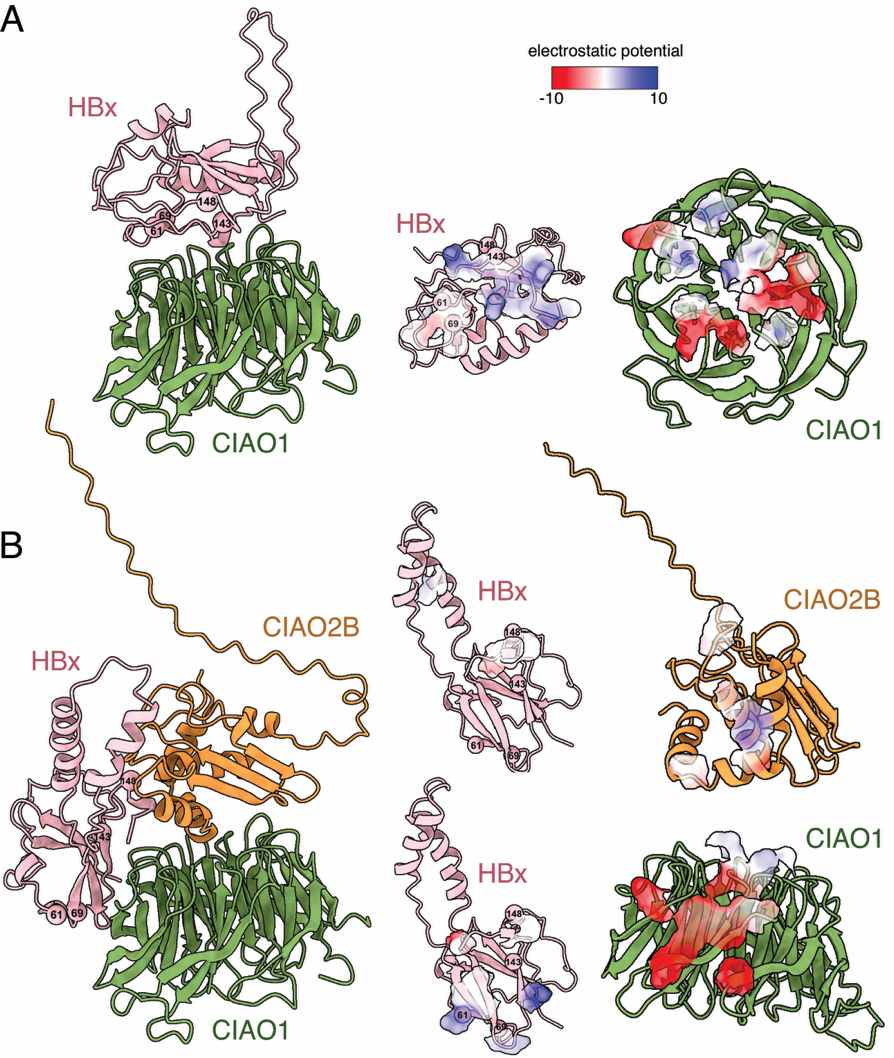

**Fig. S10. HBx potentially interacts with CIAO1 and CIAO2B through electrostatic complementarity.** AlphaFold predicted models of the CIAO1:HBx (**A**) and CIAO1:CIAO2B:HBx (**B**) complexes. Electrostatic surface potentials are mapped onto the models to highlight the predicted binding interfaces between HBx and CIAO1, and between HBx and CIAO2B. Contact residues at the binding interfaces are within an atomic distance of 3.5 Å.

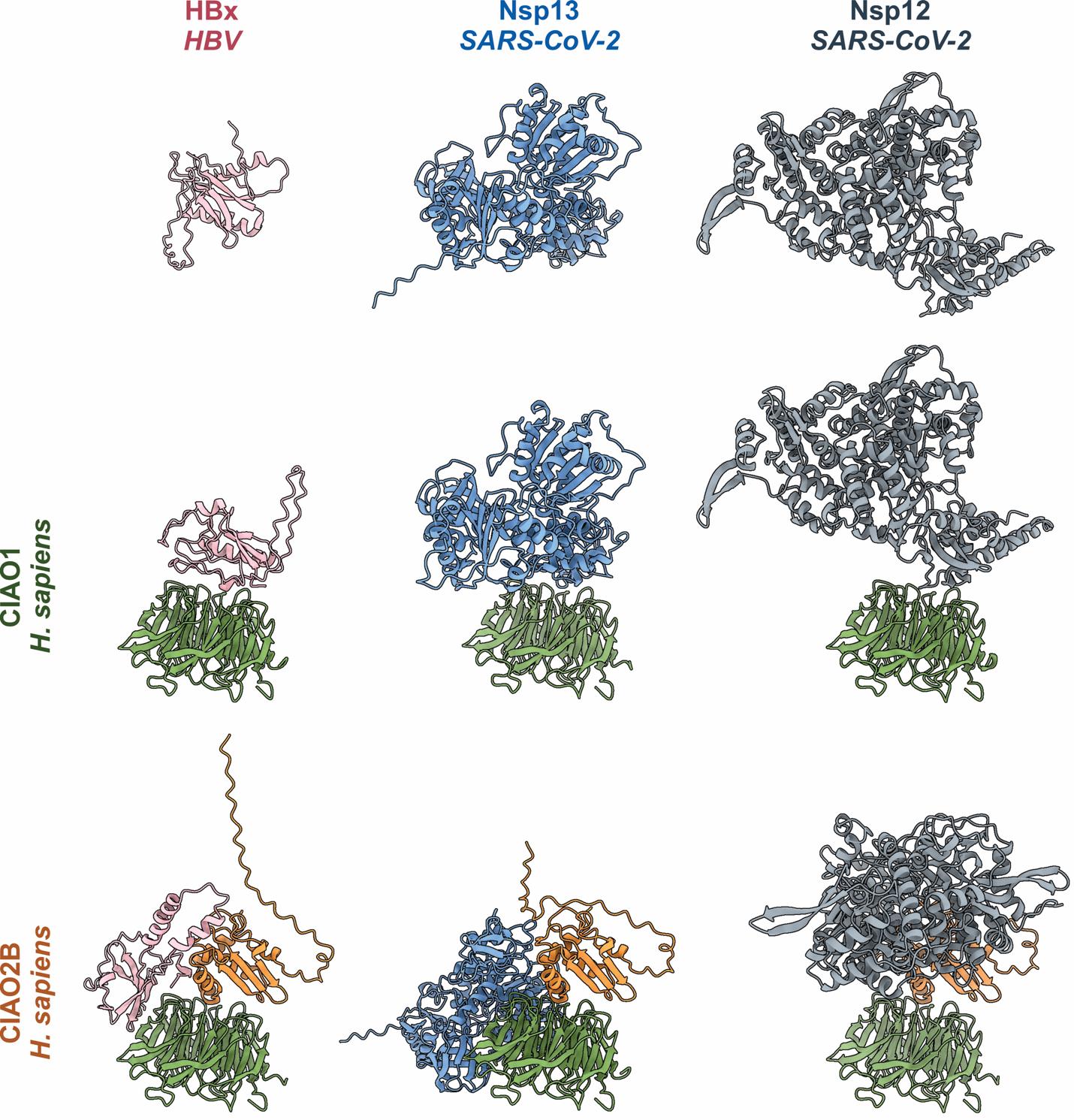

**Fig. S11.** **Structural comparison of viral proteins bound to human** **cytosolic Fe–S cluster assembly proteins CIAO1 and CIAO2B.** AlphaFold models of the individual viral proteins, HBx (pink), Nsp13 (blue), and Nsp12 (grey), are shown in three configurations: alone (top row), in complex with CIAO1 (green) (middle row), and in complex with both CIAO1 and CIAO2B (orange) (bottom row).

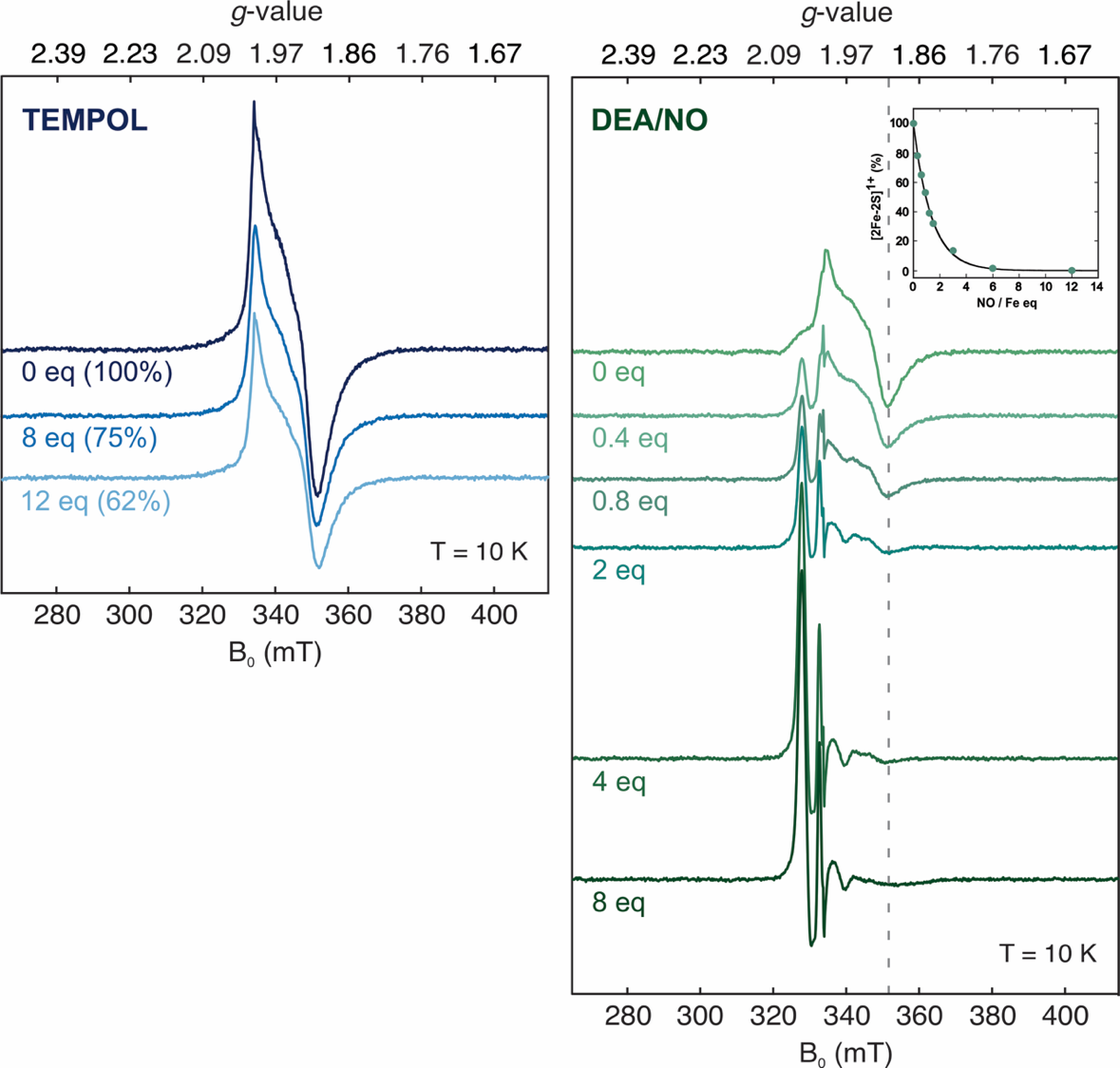

**Fig. S12. The nitroxide radical TEMPOL and the nitric oxide donor DEA/NO disassemble the [2Fe-2S] cluster of HBx.** WT DsbC-HBx was co-expressed with the pDB1282 plasmid in M9 media supplemented with Fe(NH_4_)_2_(SO_4_)_2_, then isolated under ambient conditions to obtain the [2Fe-2S]^2+^ form. The cluster was reduced with an excess of sodium dithionite (6 mM) for 30s following by rapid freezing to obtain EPR active [2Fe-2S]^1+^ form. CW EPR spectra of aerobically purified protein were recorded following incubation with increasing equivalents of TEMPOL (left) and DEA/NO (right, 1 mol DEA/NO liberates 1.5 mol NO). Experimental conditions: microwave frequency = 9.38 GHz, temperature = 10 K, microwave power = 0.64 mW, modulation amplitude = 1 mT. Quantification was performed by double integration, and the integrated intensities of the [2Fe-2S]^1+^ cluster signal were normalized to that of 0 eq TEMPOL and DEA/NO. The characteristic [2Fe-2S]^1+^ cluster trough at *g*-value = 1.90 is indicated by a dashed line. The titration curve (inset) shows the extent of [2Fe-2S]^1+^ cluster degradation as a function of NO to Fe equivalents in HBx.

**Tables**

**Table S1. LC-MS/MS results for the [4Fe-4S] cluster-containing form of HBx reconstituted with semi-enzymatic synthesis employing IscS.**

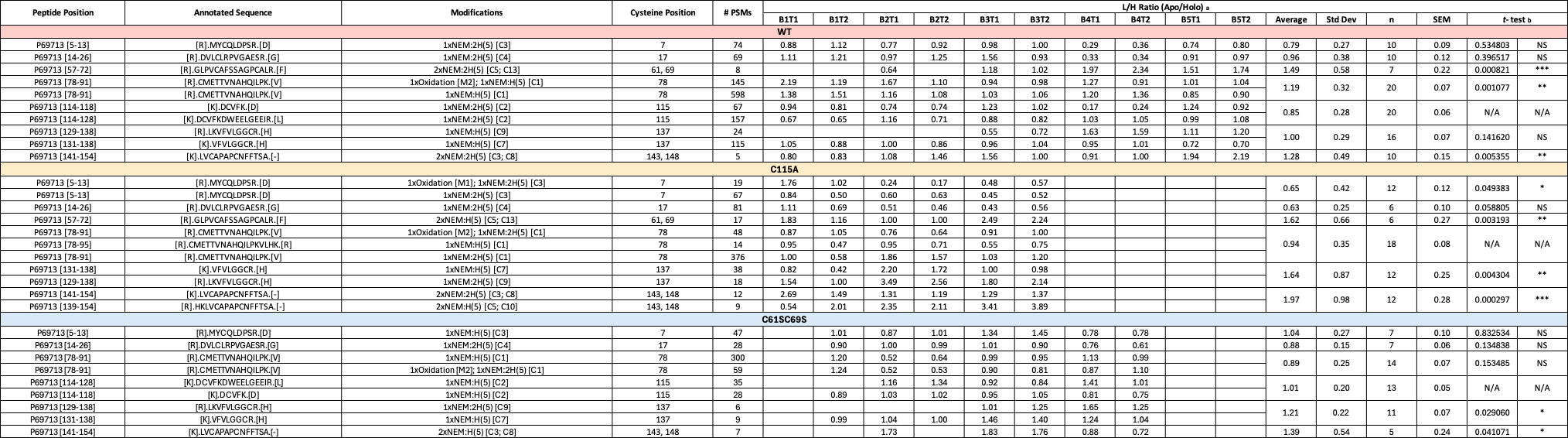

^a^ The notation BxTy refers to the y-th technical (T) replicate from the x-th biological (B) replicate.

^b^ The independent *t*-test is used to compare the mean L/H ratio value between each cysteine residue, with C115 as the control group for WT and the C61SC69S variant, and C78 as the control group for the C115A variant. Significance is calculated as NS (not significant), **p* < 0.05, ***p* < 0.01, and ****p* < 0.001.

**Table S2. LC-MS/MS results for independently expressed and isolated [4Fe-4S] and [2Fe-2S] cluster-containing forms of HBx.**

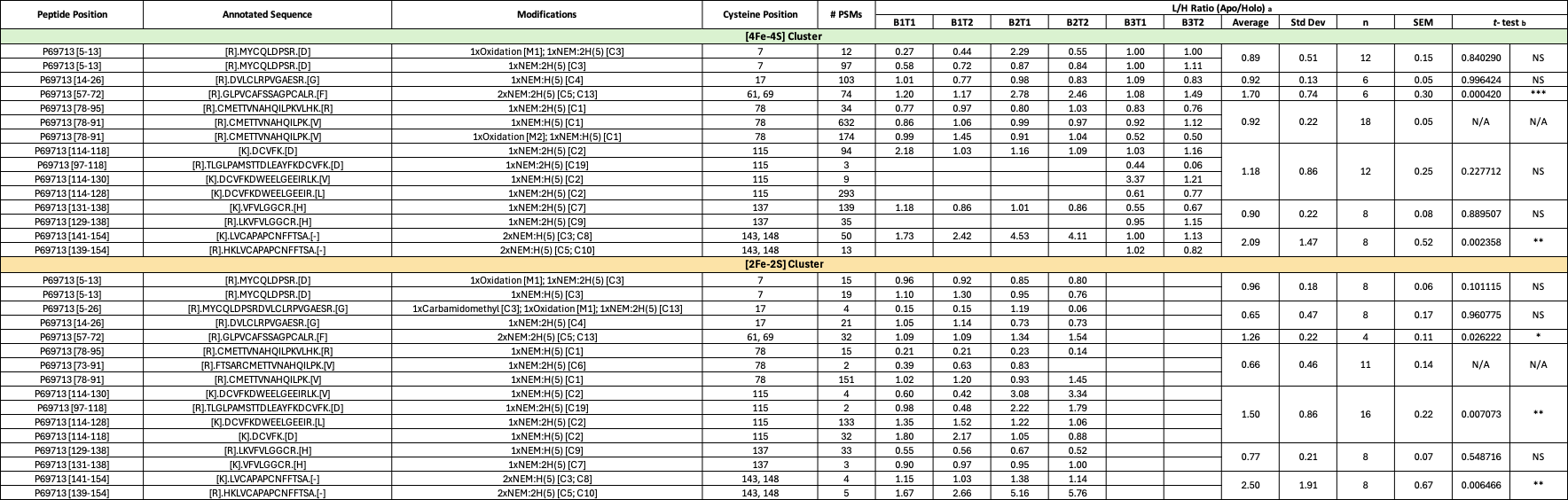

^a^ The notation BxTy refers to the y-th technical (T) replicate from the x-th biological (B) replicate.

^b^ The independent *t*-test is used to compare the mean L/H ratio value between each cysteine residue, with C78 as the control group for both WT and the C61SC69S variant. Significance is calculated as NS (not significant), **p* < 0.05, ***p* < 0.01, and ****p* < 0.001.

**Table S3. LC-MS/MS results for the Zn reconstituted HBx.**

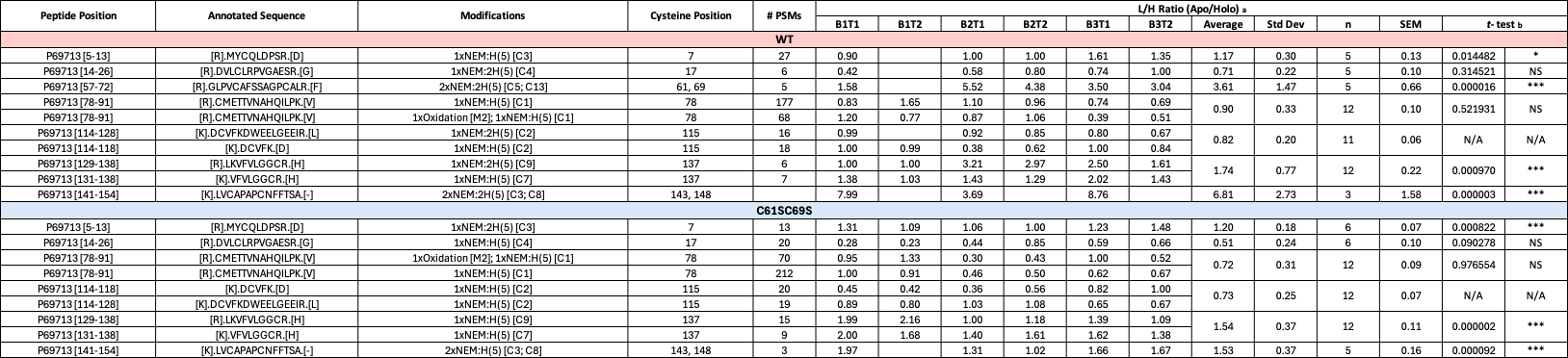

^a^ The notation BxTy refers to the y-th technical (T) replicate from the x-th biological (B) replicate.

^b^ The independent *t*-test is used to compare the mean L/H ratio value between each cysteine residue, with C115 as the control group for both WT and the C61SC69S variant. Significance is calculated as NS (not significant), **p* < 0.05, ***p* < 0.01, and ****p* < 0.001.

**Table S4. LC-MS/MS identification of proteins from excised bands** **corresponding to CIAO1, CIAO2, and HBx in the anti-His Western blot of CIAO1:CIAO2B:HBx co-expression (Fig. 5B)**

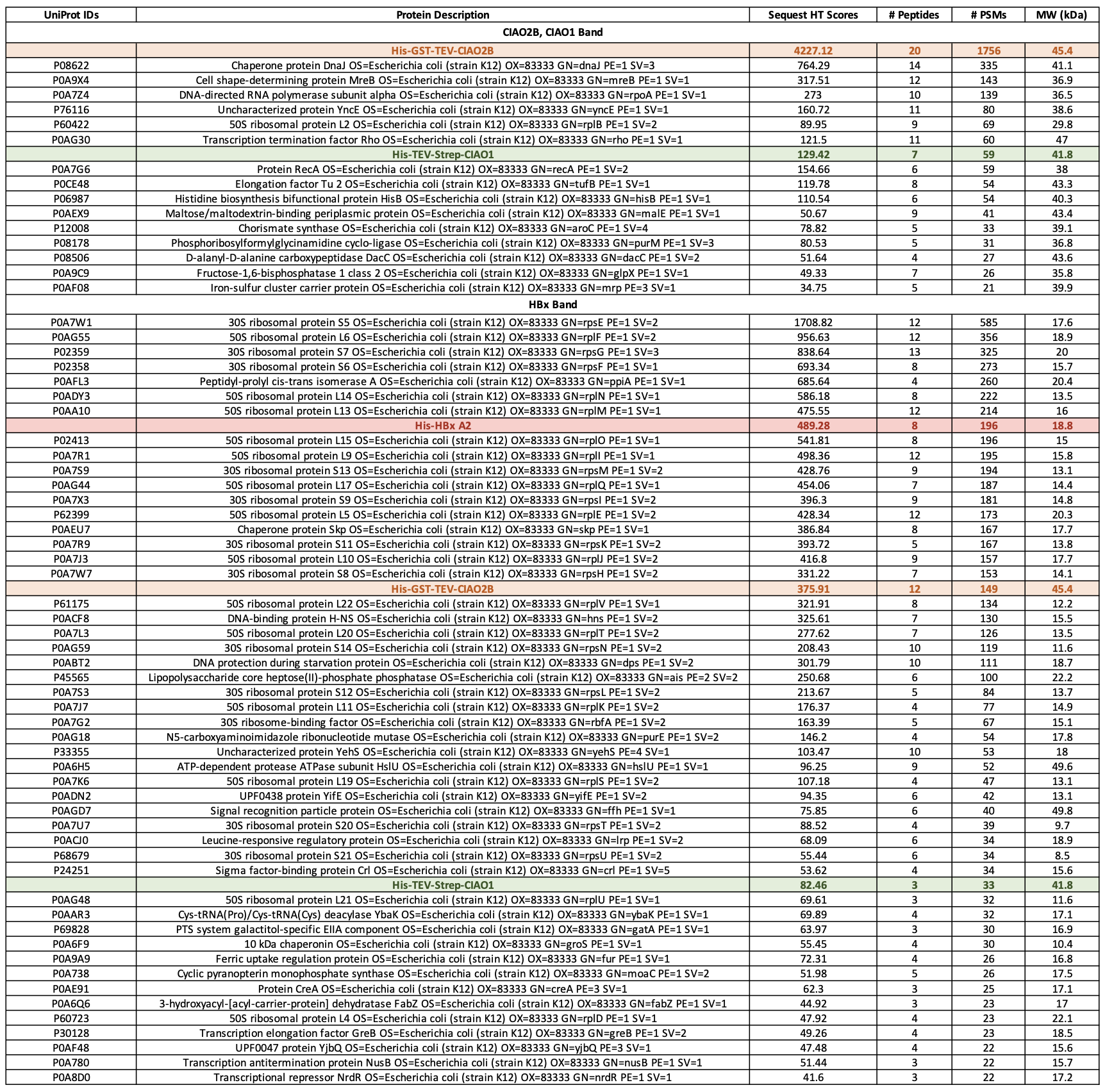
